# Exploiting a PAX3-FOXO1-induced synthetic lethal ATR dependency for rhabdomyosarcoma therapy

**DOI:** 10.1101/2020.12.04.411413

**Authors:** Heathcliff Dorado García, Yi Bei, Jennifer von Stebut, Glorymar Ibáñez, Koshi Imami, Dennis Gürgen, Jana Rolff, Konstantin Helmsauer, Natalie Timme, Victor Bardinet, Rocío Chamorro González, Ian C. MacArthur, Fabian Pusch, Celine Y. Chen, Joachim Schulz, Antje M. Wengner, Christian Furth, Birgit Lala, Angelika Eggert, Georg Seifert, Patrick Hundsoerfer, Marieluise Kirchner, Philipp Mertins, Matthias Selbach, Andrej Lissat, Johannes H. Schulte, Kerstin Haase, Monika Scheer, Michael V. Ortiz, Anton G. Henssen

**Affiliations:** Experimental and Clinical Research Center (ECRC) of the MDC and Charité Berlin, Berlin, Germany; Department of Pediatric Oncology and Hematology, Charité – Universitätsmedizin Berlin, corporate member of Freie Universität Berlin, Humboldt-Universität zu Berlin, Berlin, Germany; Department of Pediatrics, Memorial Sloan Kettering Cancer Center, New York City, NY, USA; Max Delbrück Center for Molecular Medicine, Berlin, Germany; Experimental Pharmacology and Oncology (EPO), Berlin, Germany; Bayer AG, Berlin, Germany; Berlin Institute of Health, 10178 Berlin, Germany; German Cancer Consortium (DKTK), partner site Berlin, and German Cancer Research Center (DKFZ), Heidelberg, Germany

## Abstract

Pathognomonic PAX3-FOXO1 fusion oncogene expression is associated with poor outcome in rhabdomyosarcoma. Combining genome-wide CRISPR screening with cell-based functional genetic approaches, we here provide evidence that PAX3-FOXO1 induces replication stress, resulting in a synthetic lethal dependency to ATR-mediated DNA damage-response signaling in rhabdomyosarcoma. Expression of PAX3-FOXO1 in muscle progenitor cells was not only sufficient to induce hypersensitivity to ATR inhibition, but PAX3-FOXO1-expressing rhabdomyosarcoma cells also exhibited increased sensitivity to structurally diverse inhibitors of ATR, a dependency that could be validated genetically. Mechanistically, ATR inhibition led to replication stress exacerbation, decreased BRCA1 phosphorylation and reduced homologous recombination-mediated DNA repair pathway activity. Consequently, ATR inhibitor treatment increased sensitivity of rhabdomyosarcoma cells to PARP inhibition *in vitro*, and combined ATR and PARP inhibition induced regression of primary patient-derived alveolar rhabdomyosarcoma xenografts *in vivo*. Moreover, a genome-wide CRISPR activation screen (CRISPRa) identified *FOS* gene family members as inducers of resistance against ATR inhibitors. Mechanistically, *FOS* gene family members reduced replication stress in rhabdomyosarcoma cells. Lastly, compassionate use of ATR inhibitors in two pediatric patients suffering from relapsed PAX3-FOXO1-expressing alveolar rhabdomyosarcoma showed signs of tolerability, paving the way to clinically exploit this novel synthetic lethal dependency in rhabdomyosarcoma.

## Introduction

Rhabdomyosarcomas are the most common soft tissue tumors in childhood^1^. Whereas the majority of rhabdomyosarcomas are histologically classified as embryonal rhabdomyosarcoma (ERMS), about 25% of cases present as alveolar rhabdomyosarcoma (ARMS) and harbor pathognomonic chromosomal translocations involving genes encoding for the *PAX3* (and less frequently, *PAX7*) and *FOXO1* transcription factors^2,3^. PAX3/7-FOXO1 expression is not only sufficient to drive tumorigenesis^4^, but is also associated with adverse clinical outcome. Patients harboring PAX3/7-FOXO1-expressing rhabdomyosarcomas often develop metastases and/or resistance to chemotherapy during the course of disease^5^. Despite advances in targeted cancer therapy development, no new pharmacological treatment options have been clinically approved for metastatic or recurrent rhabdomyosarcomas in the last ∼30 years^6^. It is widely accepted that current treatment strategies have reached their limits. To overcome these limits, new therapeutically actionable disease mechanisms need to be identified. However, PAX3/7- FOXO1-driven rhabdomyosarcomas are rarely associated with therapeutically actionable genetic aberrations^7^. Thus, the identification of new therapeutic strategies for high-risk PAX3/7-FOXO1-expressing rhabdomyosarcoma remains challenging.

Most cancers depend on active DNA damage repair, explaining why genotoxic agents are among the most effective chemotherapeutic agents in cancer therapy^8^. The therapeutic window of genotoxic agents, however, is often narrow and considerable long-term sequelae occur in patients treated with such agents. Synthetic lethal cellular dependencies have emerged as tumor-specific vulnerabilities, which provide therapeutic targets offering much broader therapeutic windows^9^. In particular, DNA damage response (DDR) pathway dependencies are being successfully exploited for the development of novel therapies. As a prototypical example, *BRCA1* deficient tumors rely on PARP- mediated base-excision DNA repair (BER), a synthetic lethal relationship that was clinically translated in breast and ovarian cancers among other tumor entities^10,11^. Thus, exploiting DDR pathway dependencies may enable the development of novel therapeutic strategies for rhabdomyosarcomas.

Oncogenes, particularly those encoding for transcription factors and fusion transcription factors, can interfere with the normal function of the DNA replication machinery through massive deregulation of transcriptional activity^12^. Resulting transcription-induced replication fork stalling leads to activation of DDR pathways, during which unprotected single stranded DNA is bound by Replication Protein A (RPA32), subsequently recruiting the ataxia telangiectasia and Rad3 related (ATR) kinase^13-16^. This process has been termed oncogene-induced replication stress. Upon recognition of the DNA break, ATR activates checkpoint kinase 1 (CHK1) among other factors to stop cells from cycling and to coordinate DNA repair^17^. Unsurprisingly, many tumors depend on ATR activity to proliferate in the presence of oncogene-induced replication stress. Based on this observation, ATR has become a candidate target for pharmacological inhibition in cancer therapy. Even though some molecular features create synthetic lethal ATR dependencies, including *ATM* and *TP53* loss, MYC proto-oncogene expression, EWS-FLI1 fusion oncogene expression and PGBD5 expression^18-26^, the mechanisms driving response to ATR inhibition in most cancer entities are still largely unknown. Here, we show that PAX3-FOXO1 expression renders highly aggressive alveolar rhabdomyosarcomas sensitive to ATR inhibition *in vitro* and *in vivo*, revealing a novel and clinically exploitable synthetic lethal dependency in rhabdomyosarcoma.

## Results

### Alveolar rhabdomyosarcoma cell lines genetically depend on ATR and are hypersensitive to pharmacological ATR pathway inhibition

To identify therapeutically actionable DNA damage repair pathway vulnerabilities in alveolar rhabdomyosarcomas, we inquired the publicly available database, DepMap^27^, focusing on pharmacologically targetable DDR pathway members ATR, ATM, CHK1 and CHK2 (Fig. 1a and Extended data Fig. 1a-c). Whereas ATM and CHK2 depletion via CRISPR-Cas9 did not lead to significant survival disadvantage in rhabdomyosarcoma cell lines, depletion of ATR and CHK1 was strongly selected against, as evidenced by low dependency scores (Fig. 1a and Extended data Fig. 1a). Intriguingly, alveolar rhabdomyosarcoma cells were amongst the most dependent on ATR (Fig. 1a). To test if this genetic dependency could translate into sensitivity towards pharmacological ATR inhibition, we incubated a panel of nine rhabdomyosarcoma cell lines, all of which expressed ATR and CHK1 (Extended data Fig. 1d-e), with structurally diverse ATR- (AZD6738, BAY-1895344, VE-822), ATM- (AZD0156, KU60019) and CHK1/2- specific (AZD7762) small molecule inhibitors (Fig. 1b-e and Extended data Fig. 2a-h). Rhabdomyosarcoma cell lines showed varying degrees of sensitivity to small molecule-mediated ATR, ATM and CHK1/2 inhibition, with inhibitory concentrations (IC_50_) ranging between 50 nM and 10 µM (Extended data Fig. 2i). In line with greater genetic dependency towards ATR (Fig. 1a), alveolar rhabdomyosarcoma cell lines were significantly more sensitive to ATR inhibition by three structurally diverse small molecule ATR inhibitors AZD6738, VE-822 and BAY-1895344 than embryonal rhabdomyosarcoma cells (Fig. 1b-e and Extended data Fig. 2a-b). This increased sensitivity was also observed for CHK1/2 inhibitor AZD7762 (Extended data Fig. 2c-d), indicating that alveolar rhabdomyosarcoma cells depend on active ATR pathway signaling. In line with an ATR pathway-specific dependency, no difference in sensitivity between alveolar and embryonal rhabdomyosarcoma cells was observed for ATM inhibitors KU60019 and AZD0156 (Extended data Fig. 2e-h). Thus, alveolar rhabdomyosarcoma cells depend on ATR, rendering them hypersensitive to pharmacological ATR pathway inhibition.

**Fig. 1.**
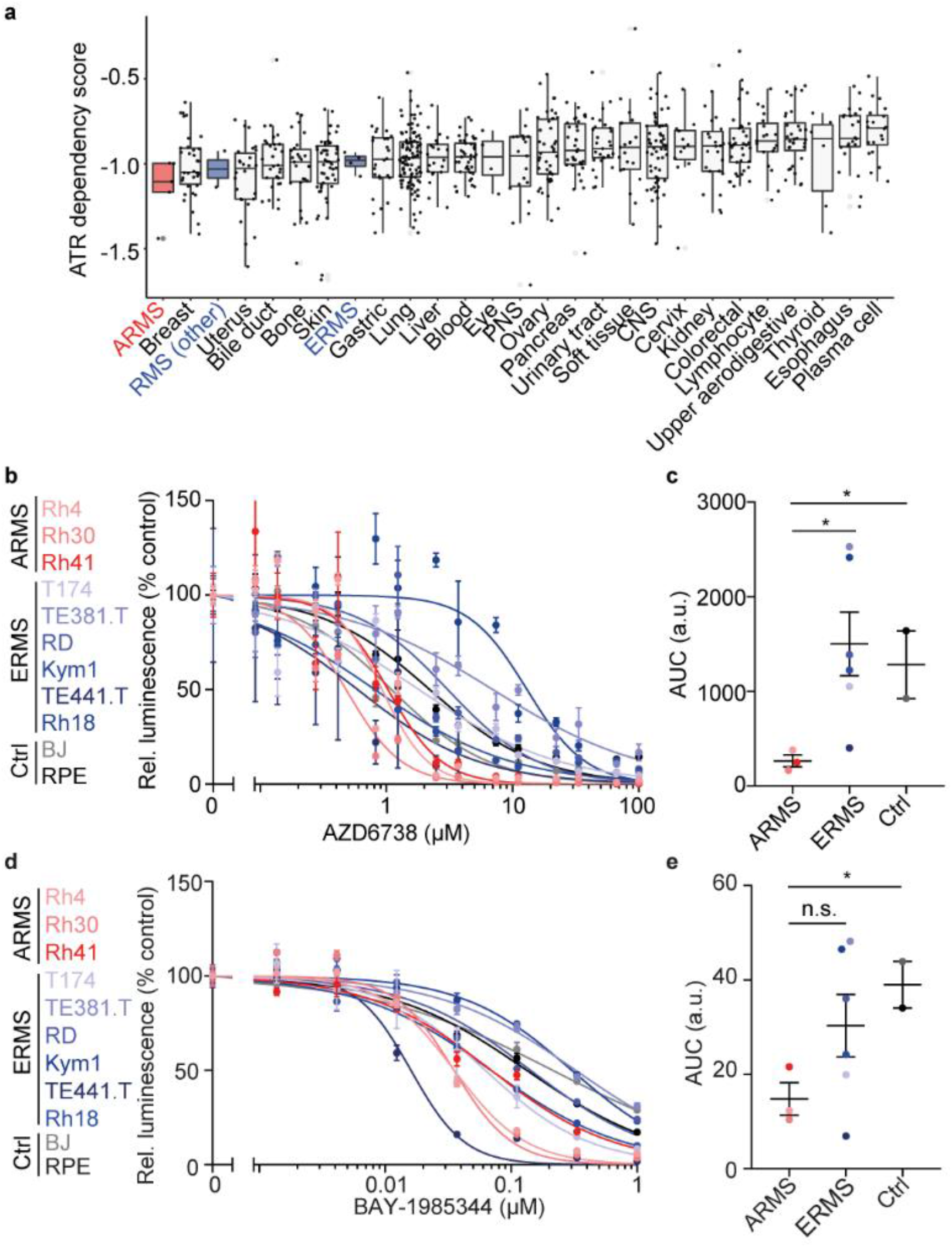
Alveolar rhabdomyosarcoma cells depend on ATR and are hypersensitive to pharmacological ATR inhibition. **(a)** ATR dependency scores in cell lines from a variety of cancer entities obtained from CRISPR (Avana) 20Q2 (Alveolar rhabdomysarcoma, ARMS, red; other rhabdomyosarcoma and embryonal rhabdomyosarcoma, ERMS, blue). (**b)** Dose-response curves of cell viability for ARMS cell lines (red) and ERMS (blue) treated with the ATR inhibitor AZD6738 compared to non-transformed cells (grey; *n*=3). **(c)** Area under the curve (AUC) of dose-response curves shown in (b) (*P*=0.041 for ARMS vs ERMS; *P*=0.035 for ARMS vs Ctrl). **(d)** Dose-response curves of cell viability for ARMS cell lines (red) and ERMS (blue) treated with the ATR inhibitor BAY-1895344 compared to non-transformed cells (grey; *n*=3). (**e)** AUC the dose-response curves shown in (c) (*P*=0.162 for ARMS vs ERMS; *P*= 0.025 for ARMS vs Ctrl; two-sided student’s t-test; error bars represent standard error of the mean).

### ATR inhibition leads to replication stress, genomic instability, apoptosis and cell cycle disruption in alveolar rhabdomyosarcoma cells

ATR is a key regulator of replication stress and genomic stability^28^. We hypothesized that high steady-state replication stress and genomic instability may cause sensitivity of alveolar rhabdomyosarcoma cells to ATR inhibition. Once recruited to single stranded DNA bound by RPA32 during replication stress, ATR phosphorylates RPA32 at T21 and S33^29^. To test the association between steady-state replication stress and ATR inhibitor sensitivity, we measured RPA32 phosphorylation at T21 and S33 in rhabdomyosarcoma cell lines using western immunoblotting. Indeed, high levels of RPA32 phosphorylation were observed in alveolar rhabdomyosarcoma cell lines (Fig. 2a-b). In line with higher replication stress, a higher fraction of alveolar rhabdomyosarcoma cells presented micronuclei (Fig. 2c-d), a sign of genomic instability associated with replication stress^30^. This suggested that alveolar rhabdomyosarcoma cells either had critically high DNA damage levels or lacked intrinsic DNA damage repair activity to resolve such damage. To distinguish between these possible mechanisms, we assessed the endogenous activity of two major DNA damage repair mechanisms, non-homologous end-joining repair (NHEJ) and homologous recombination (HR), across rhabdomyosarcoma cell lines. No significant difference in NHEJ and HR activity was observed between alveolar and embryonal rhabdomyosarcoma cell lines (Fig. 2e and Extended data Fig. 3a-d), suggesting that differences in endogenous DNA damage levels rather than repair activity may drive the observed differences in steady-state genomic instability.

**Fig. 2.**
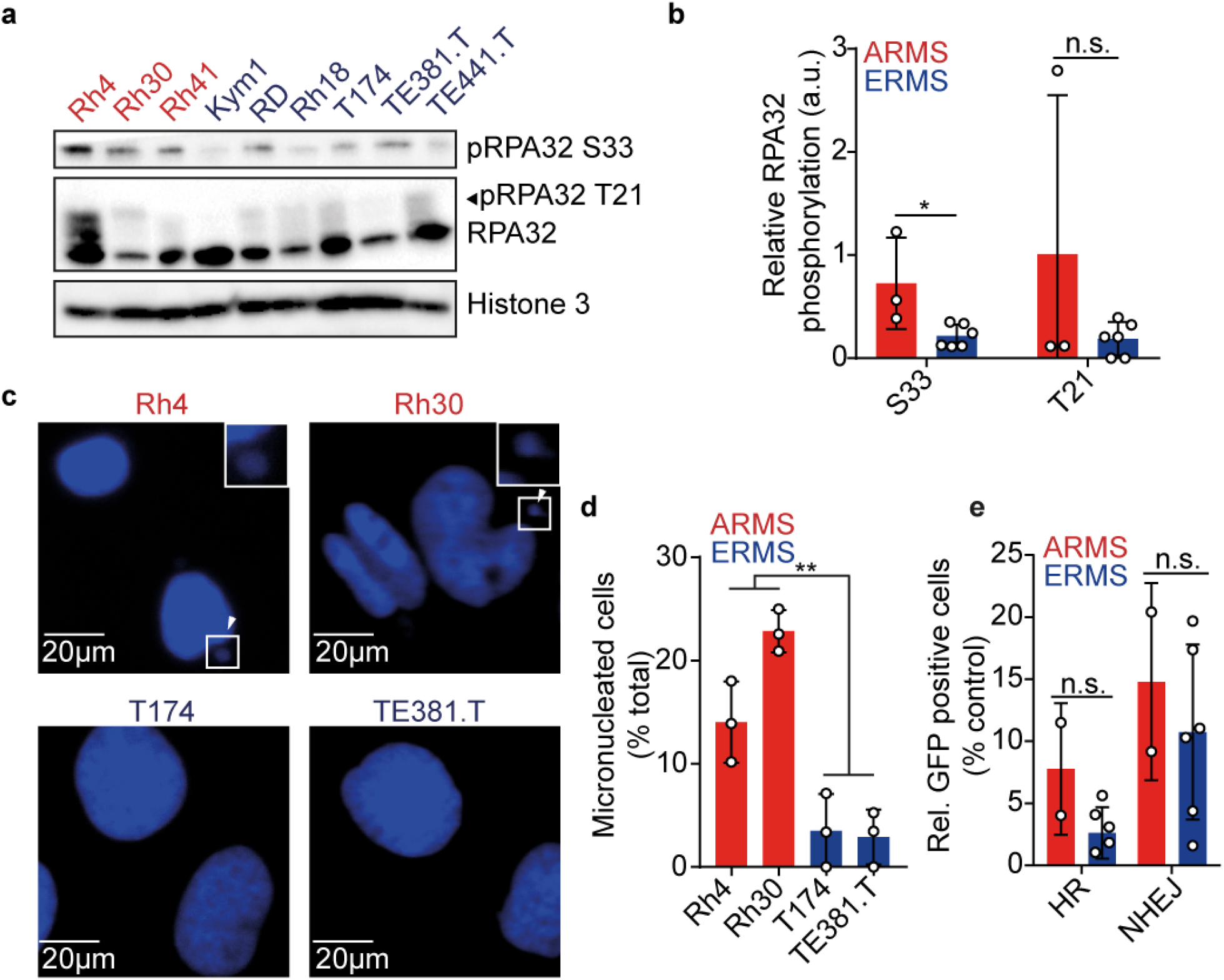
Alveolar rhabdomyosarcoma cells present higher baseline genomic instability than embryonal rhabdomyosarcoma despite similar DNA repair efficiencies. **(a)** Western immunoblot for RPA32 phosphorylation in rhabdomyosarcoma cell lines (Histone 3 serves as a loading control). **(b)** Quantification of RPA32 S33 (*P*=0.025) and T21 (*P*=0.208) signal intensity. **(c)** Representative images of rhabdomyosarcoma cells stained with DAPI (arrows indicate micronuclei, squares in top right corners indicate higher magnification). **(d)** Quantification of the fraction of rhabdomyosarcoma cells containing at least one micronucleus (*P*=1.4×10^−4^). **(e)** Quantification of HR (P=0.071) and NHEJ activity (P=0.516) in rhabdomyosarcoma cell lines (two-sided student’s t-test; error bars represent standard error of the mean).

We hypothesized that ATR inhibition may lead to exacerbation of replication stress and genomic instability, i.e. an increase in the severity above a tolerable point, specifically in alveolar rhabdomyosarcoma cells. Indeed, RPA32 S33 and T21 phosphorylation increased after treatment with AZD6738 in all cell lines tested (Fig. 3a and Extended data Fig. 3e-f) and was accompanied by an increase in micronucleated cells (Fig. 3b and Extended data Fig. 3g), in particular in alveolar rhabdomyosarcoma cells. Co-staining with 5-Ethynyl-2′-deoxyuridine (EdU) and propidium iodide (PI) after incubation with ATR inhibitors revealed that alveolar rhabdomyosarcoma cells accumulated in G2/M-phase with a corresponding reduction of cells in S-phase, indicating a bypass of intra-S phase cell cycle checkpoint (Fig. 3c and Extended data Fig. 3g). In contrast, embryonal rhabdomyosarcoma cells’ cycling was almost unperturbed (Fig. 3c). Consistent with the increased ATR dependency (Fig. 1a), alveolar rhabdomyosarcoma cells showed significant accumulation of unrepaired DNA double stranded breaks when treated with an ATR inhibitor, as measured by terminal deoxynucleotidyl transferase dUTP nick end labeling (TUNEL; Fig. 3d and Extended data Fig. 3h). Furthermore, increased cell death, as measured by Caspase 3 cleavage, was more pronounced in alveolar compared to embryonal rhabdomyosarcoma cells incubated in the presence of an ATR inhibitor (Fig. 3e and Extended data Fig. 3i). Lastly, the fraction of aneuploid cells significantly increased after ATR inhibition (Fig. 3f and Extended data Fig. 3j), suggesting chromosome missegregation due to erroneous repair of unresolved replication intermediates. Based on this, we propose that ATR inhibition exacerbates replication stress, in particular in alveolar rhabdomyosarcoma cells, which enter mitosis with unrepaired DNA damage, leading to irreparable DNA damage during mitosis incompatible with cell survival.

**Fig. 3.**
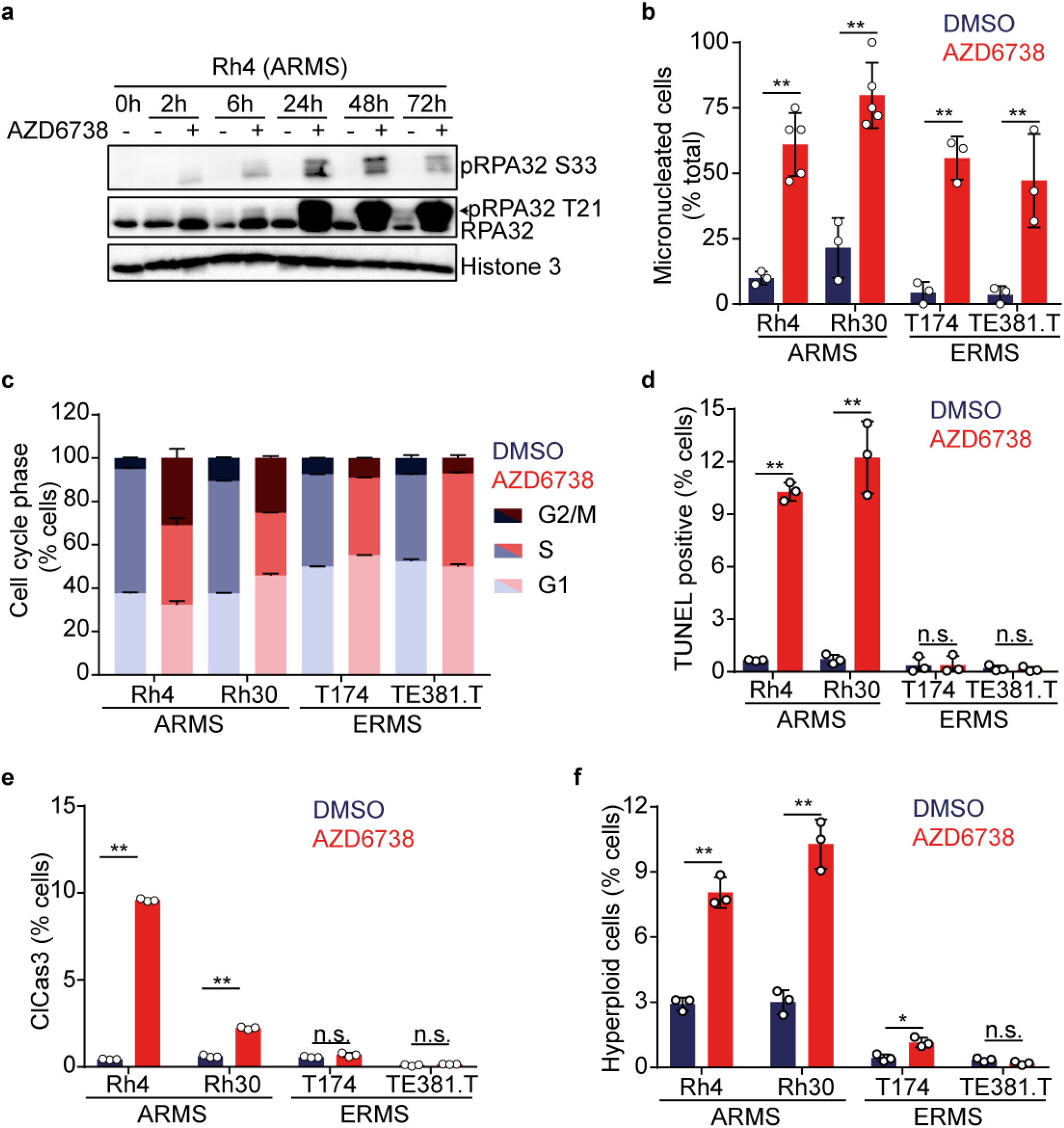
Pharmacological ATR inhibition exacerbates replication stress and leads to genomic instability, apoptosis and cell cycle disruption in alveolar rhabdomyosarcoma cells. **(a)** Western immunoblot of RPA32 phosphorylation in Rh4 cells treated with ATR inhibitor AZD6738 (750 nM). **(b)** Fraction of micronucleated cells after incubation with AZD6738 (750 nM) for 72 hours (*n*=3, with at least 40 nuclei counted per condition; *n*=5, for samples with less than 40 nuclei per image; from left to right, *P*=7.1×10^−4^, *P*=4.6×10^−4^, *P*=1.7×10^−4^ and *P*=0.009). **(c)** Cell cycle phase distribution as measured using flow cytometry analysis of EdU and propidium iodide-labeled rhabdomyosarcoma cells treated with AZD6738 compared to DMSO-treated cells (*n*=3). **(d)** Unrepaired double strand breaks (DSBs) measured by TUNEL in cells treated with AZD6738 (750 nM) for 72 hours (*n*=3; from left to right, *P*=6.0×10^−6^, *P*=6.5×10^−6^, *P*=0.489 and *P*=0.450). **(e)** Fraction of apoptotic cells treated with AZD6738 (750 nM), as measured by cleaved caspase 3 staining (*n*=3; from left to right, *P*=2.3×10^−9^, *P*=5.4×10^− 6^, *P*=0.062 and *P*=0.054). **(f)** Fraction of hyperploid/aneuploid rhabdomyosarcoma cells treated with AZD6738 (750 nM) (*n*=3; from left to right, *P*=3.2×10^−4^, *P*=5.6×10^−5^, *P*=0.010 and *P*=0.080; two-sided student’s t-test; error bars represent standard error of the mean).

### PAX3-FOXO1 is sufficient to increase replication stress and is required for hypersensitivity to ATR pathway inhibition

Several factors exist in synthetic lethal relationship with ATR^18-26^. To identify which of the known factors were responsible for sensitivity to ATR inhibition in alveolar rhabdomyosarcoma cells, we assessed their presence in rhabdomyosarcoma cell lines and their association with ATR inhibitor sensitivity (Extended data Fig. 4). Even though some cell lines that were highly sensitive to ATR inhibition also presented reduced expression of TP53 or ATM, or high *PGBD5* mRNA expression, these associations were not statistically significant (Extended data Fig. 4a-b and Fig. 4d-f). In line with MYCN’s ability to drive replication stress, high MYCN expression, as measured using western immunoblotting, was significantly associated with high ATR inhibitor sensitivity both in alveolar and embryonal rhabdomyosarcoma cells (Extended data Fig. 4a and Fig. 4c). Intriguingly, the pathognomonic fusion oncogene PAX3-FOXO1 drives MYCN expression in alveolar rhabdomyosarcomas^31^. Based on this function of PAX3-FOXO1 and previous reports showing that chimeric transcription factors, such as EWS-FLI1 in Ewing sarcoma, can themselves render cells hypersensitive to ATR inhibition through induction of replication stress^32,33^, we hypothesized that PAX3-FOXO1 may contribute to replication stress and hypersensitivity to ATR inhibition in alveolar rhabdomyosarcomas. To test this, we ectopically expressed PAX3-FOXO1 in untransformed mouse myoblast cells (C2C12, Fig. 4a), and three embryonal rhabdomyosarcoma cell lines (Extended data Fig. 5a-c). PAX3-FOXO1 expression sensitized cells to AZD6738 and BAY-1895344, which was most pronounced in C2C12 cells (Fig. 4b and Extended data Fig. 5d-g). In line with previous reports, PAX3-FOXO1 expression in rhabdomyosarcoma cells was sufficient to induce high MYCN expression (Extended data Fig. 5a-c). Interestingly, ectopic PAX3-FOXO1 expression was associated with increased phosphorylation of RPA32 at T21, but reduced phosphorylation at S33 (Fig. 4a and Extended data Fig. 5a-c). This change in RPA32 phosphorylation was also observed in cells treated with hydroxyurea (HU), a potent inducer of replication stress, but was not observed when cells were treated with doxorubicin (Doxor.), which leads to replication stress-independent DNA damage (Fig. 4a and Extended data Fig. 5a- c). This differential effect on RPA32 phosphorylation might also be a secondary result due to cell cycle blockage in G1, as RPA32 phosphorylation at S33 is cell-cycle dependent and accumulates in late S/G2^29^. Consistent with increased replication stress-induced DNA damage, H2AX phosphorylation increased in cells expressing PAX3- FOXO1 (Fig. 4c-d). To test whether PAX3-FOXO1 was required for ATR inhibitor hypersensitivity, we inducibly expressed shRNA directed against *PAX3-FOXO1* mRNA in PAX3-FOXO1-expressing alveolar rhabdomyosarcoma cells. Indeed, shRNA-mediated repression of PAX3-FOXO1 resulted in reduced sensitivity towards ATR inhibitor treatment (Fig. 4e-f), which was accompanied by reduced MYCN expression. This indicates that the pathognomonic fusion oncoprotein PAX3-FOXO1 is necessary and sufficient for ATR inhibitor sensitivity in alveolar rhabdomyosarcoma cells (Fig. 4e-f), representing a therapeutically targetable synthetic lethal ATR dependency.

**Fig. 4.**
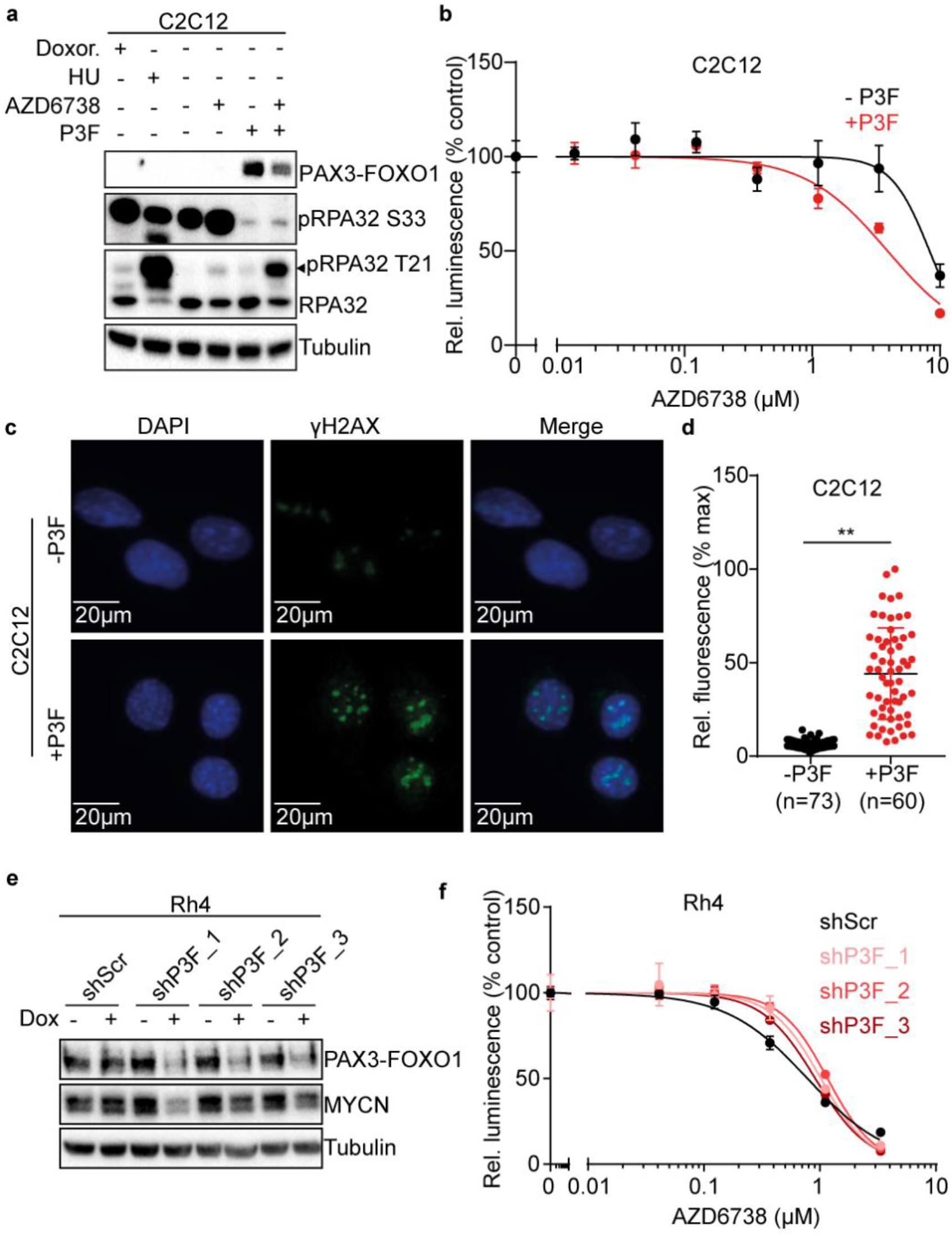
PAX3-FOXO1 is sufficient and required for hypersensitivity to ATR inhibition in myoblast cells. **(a)** Western immunoblot of PAX3-FOXO1, RPA T21 and S33 phosphorylation in C2C12 after induction of PAX3-FOXO1 expression with doxycycline (1000 ng/ml for 48 hours) and treatment with AZD6738 (750nM) compared to doxorubicin (doxor.) and hydroxyurea (HU) treated cells. **(b)** Dose-response curves of cell viability for C2C12 cells after ectopic expression of PAX3-FOXO1 and incubation with AZD6738 (*n*=3). **(c)** Representative images of H2AX phosphorylation in C2C12 cells after ectopic expression of PAX3-FOXO1, as measured using immunofluorescence. **(d)** Quantification of the H2AX phosphorylation immunofluorescence signal in C2C12 cells after ectopic expression of PAX3-FOXO1 (*P*=9.57×10^−26^). **(e)** Western immunoblot of PAX3-FOXO1 and MYCN in Rh4 cells after doxycycline-induced (1000ng/mL) expression of shRNAs targeting PAX3-FOXO1 compared to scrambled shRNA control for 48h. **(f)** Dose-response curves for Rh4 after doxycycline-induced (1000ng/mL) expression of shRNAs targeting PAX3-FOXO1 compared to scrambled shRNA control and treated with AZD6738 (*n*=3; two-sided student’s t-test; error bars represent standard error of the mean).

### A genome-wide CRISPR activation screen identifies molecular factors required for ATR inhibitor hypersensitivity in alveolar rhabdomyosarcoma cells

Successful clinical translation of synthetic lethal dependencies depends on knowledge about molecular factors influencing sensitivity to targeted therapy. Therefore, we aimed to identify factors affecting sensitivity of alveolar rhabdomyosarcoma cells to ATR inhibition, even in the presence of PAX3-FOXO1. To identify such factors, we used a genome-wide CRISPR-Cas9-based gene activation screen (CRISPRa) targeting over 70,000 genomic loci and 20,000 genes^34^. PAX3-FOXO1-expressing cells were genetically engineered to express endonuclease deficient Cas9, transcriptional activation complex members and transduced with a single guide RNA (sgRNA) library, as described before^34^. Next, cells were incubated for 9 days in the presence of 750 nM AZD6738 (Fig. 5a). SgRNAs significantly depleted in cells incubated in the presence of AZD6738 (i.e. predicted to induce a negative selective advantage under therapy) were enriched for factors promoting cell cycle checkpoint release and replication stress, e.g. *MYC* (Extended data Fig. 6a), in line with the observed PAX3-FOXO1-induced MYCN expression and replication stress leading to ATR inhibitor sensitivity (Fig. 1E), and consistent with reports in other tumor entities^22^. SgRNAs whose abundance increased in the presence of AZD6738 (i.e. predicted to induce a positive selective advantage under therapy), on the other hand, were enriched for *FOS* gene family members *FOSB, FOSL1*, and *FOSL2* (Fig. 5b-c). Efficient induction of *FOSB, FOSL1* and *FOSL2* mRNA expression by single sgRNAs was confirmed using RT-qPCR (Fig. 5d-f). Cells expressing *FOSB, FOSL1* and *FOSL2*-targeting sgRNAs and dCas9 showed reduced sensitivity to AZD6738 (Fig. 5g-h and Extended data Fig. 6b-c), as evidenced by changes in dose-response behavior of cells (Fig. 5g-h), confirming that positive enrichment in our CRISPRa screen was due to changes in sensitivity to ATR inhibition. AP-1, a transcription complex between *JNK* and *FOS* gene family members, is a known modulator of DNA damage response^35,36^, leading us to hypothesize that *FOSB, FOSL1* and *FOSL2* may reduce steady-state replication stress in alveolar rhabdomyosarcoma cells even in the presence of PAX3-FOXO1 and MYCN. Indeed, CRISPRa-driven *FOS* gene family member expression was sufficient to reduce steady state RPA S33 and T21 phosphorylation in alveolar rhabdomyosarcoma cells, indicating reduced replication stress (Fig. 5i-j). Thus, *FOS* gene family members can reduce replication stress, which is associated with decreased sensitivity of alveolar rhabdomyosarcoma cells to ATR inhibitor treatment.

**Fig. 5.**
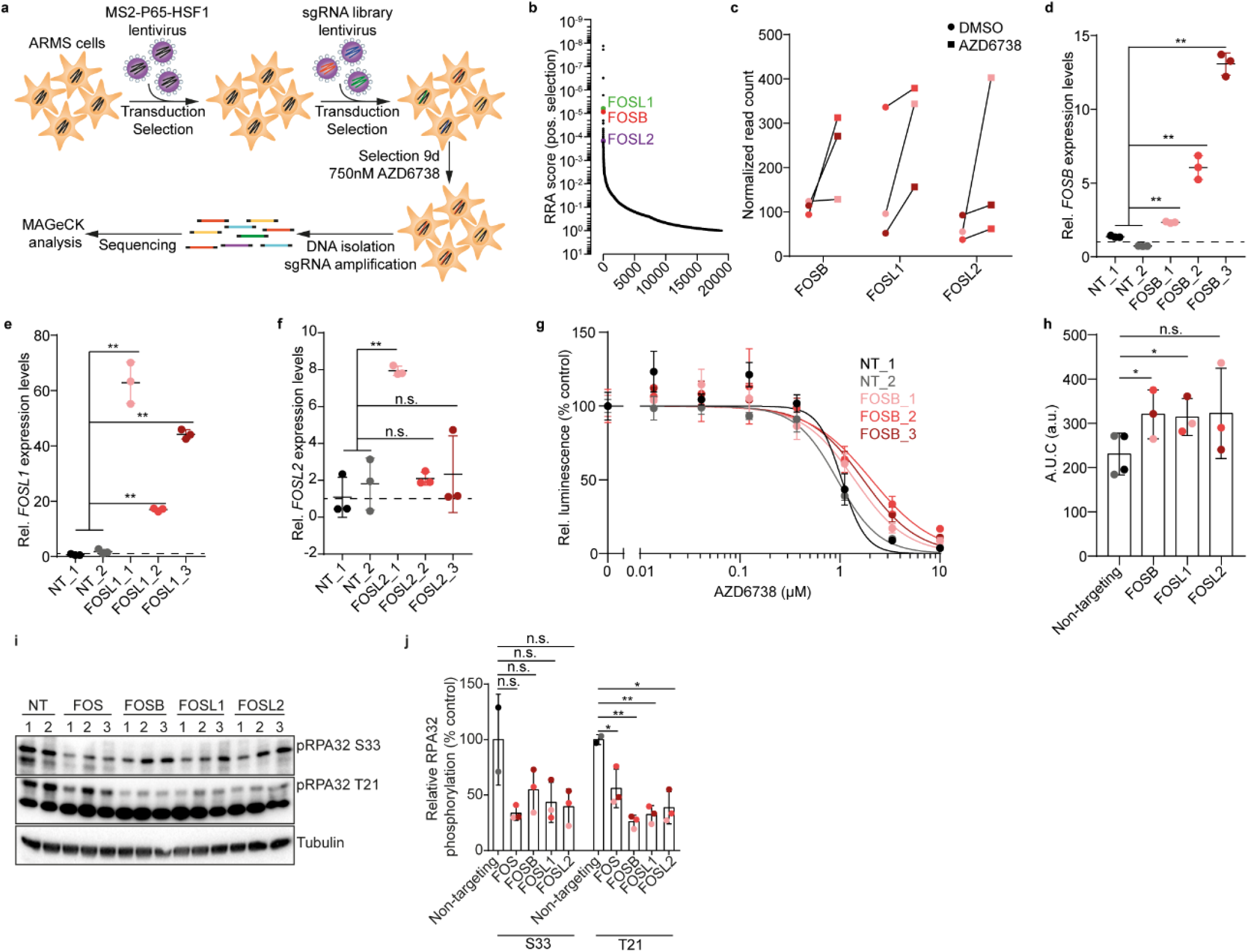
A genome-wide CRISPR-based activation screen identifies molecular modifiers of sensitivity to ATR inhibition in PAX3-FOXO1-expressing alveolar rhabdomyosarcoma cells. **(a)** Schematic representation of the genome-wide CRISPRa screen experimental design. (**b)** Waterfall plot showing the positive robust rank aggregation (RRA) score of sgRNAs in Rh4 cells incubated in the presence of AZD6738 for 9 days compared to DMSO treated cells as analyzed using MaGECK. **(c)** Relative abundance of sgRNAs targeting FOS family members *FOSB, FOSL1* and *FOSL2* in cells in the presence of AZD6738. **(d-f)** FOSB (d, *P*=4.7×10^−4^, *P*=2.9×10^−6^ and *P*=5.1×10^−9^), FOSL1 (e, *P*=1.2×10^−7^, *P*=1.6×10^−8^ and *P*=2.2×10^−10^) and FOSL2 (f, *P*=4.2×10^−5^, *P*=0.400 and *P*=0.430) mRNA expression measured using RT-qPCR in cells expressing dCas9, lentiMPH and sgRNAs targeting *FOSB, FOSL1* or *FOSL2* (*n*=3). **(g)** Representative dose-response cell viability curves of cells stably expressing dCas9, lentiMPH and sgRNAs targeting *FOSB* treated with the ATR inhibitor AZD6738. **(h)** AUC of dose-response curves in cells stably expressing dCas9, lentiMPH and sgRNAs targeting *FOSB, FOSL1* and *FOSL2* treated with the ATR inhibitor AZD6738 (*n*=3, *P*=0.029, *P*=0.034 and *P*=0.165). **(i)** Western immunoblot of RPA32 S33 and T21 phosphorylation in cells expressing sgRNAs targeting FOS family members *FOSB, FOSL1* or *FOSL2*. **(j)** Quantification of RPA32 phosphorylation compared to the corresponding non-targeting control, for S33 (*P*=0.058, 0.181, 0.113 and 0.091, respectively) and T21 (*P*=0.044, 6.43×10^−4^,1.82×10^−3^ and 0.012, respectively; two-sided student’s t-test; error bars represent standard error of the mean).

### ATR inhibition leads to reduced BRCA1 activation and repressed homologous recombination in PAX3-FOXO1-expressing rhabdomyosarcoma cells

We next sought to identify potential mechanisms of ATR inhibition-induced replication stress exacerbation and genomic instability in PAX3-FOXO1-expressing alveolar rhabdomyosarcoma cells by measuring proteome-wide changes in phosphorylation using stable isotope labeling with amino acids in cell culture (SILAC) followed by liquid chromatography with tandem mass spectrometry (LC-MS/MS) phospho-proteomic analysis of cells incubated in the presence of AZD6738. Short-term AZD6738 incubation for two hours significantly reduced phosphorylation of known ATR pathway members (Fig. 6a), such as the direct ATR target TP53 S15^37^. Using a phosphosite-centered tool^38^, we analyzed the activated and repressed pathways after pharmacological ATR inhibition (Fig. 6b). As expected, the ATR pathway was the most significantly repressed pathway, followed by the ATM and response to UV irradiation pathways. In line with the observed accumulation of cells in G2/M phases after ATR inhibition, peptides from the CDK1 pathway and pathways activated in response to nocodazole, an inhibitor of microtubule formation leading to lack of mitotic spindle and M-phase arrest^39^, were phosphorylated at higher degrees after ATR inhibitor treatment (Fig. 6b). Homologous recombination (HR) and post-replication repair were among the cellular processes most significantly repressed after ATR inhibition (Fig. 6c), supporting our conclusion that the observed increase in genomic instability in cells was due to erroneous repair of unresolved replication intermediates. A particularly high degree of differential phosphorylation was measured in BRCA1 peptides (Fig. 6a). BRCA1 is a known substrate of ATR^40^, and is involved in HR at sites of replication stress^41-43^. A cluster of BRCA1 serine residues, including S1524, can be phosphorylated by ATR and serve as key regulatory sites for BRCA1 activity in DNA damage repair^40,44,45^. Using western immunoblotting, we tracked phosphorylation of one of these residues, BRCA1 S1524, in cells over the course of AZD6738 treatment. Increased replication stress in exponentially growing cells was accompanied with increased S1524 phosphorylation, which was significantly reduced following AZD6738 treatment (Fig. 6d-e), confirming LC-MS/MS-based measurements (Fig. 6a). Whereas short-term ATR inhibition led to decreased BRCA1 S1524 phosphorylation, long-term treatment was also accompanied with decreased total BRCA1 levels (Fig. 6d-e). Next, we tested whether reduced BRCA1 phosphorylation affected BRCA1 activity during HR by measuring HR and NHEJ in cells after incubation with AZD6738. Indeed, HR activity, and to a lesser extent NHEJ activity, were significantly reduced in cells incubated with AZD6738 (Fig. 6f). Thus, ATR inhibition represses BRCA1 activity and HR in rhabdomyosarcoma cells.

**Fig. 6.**
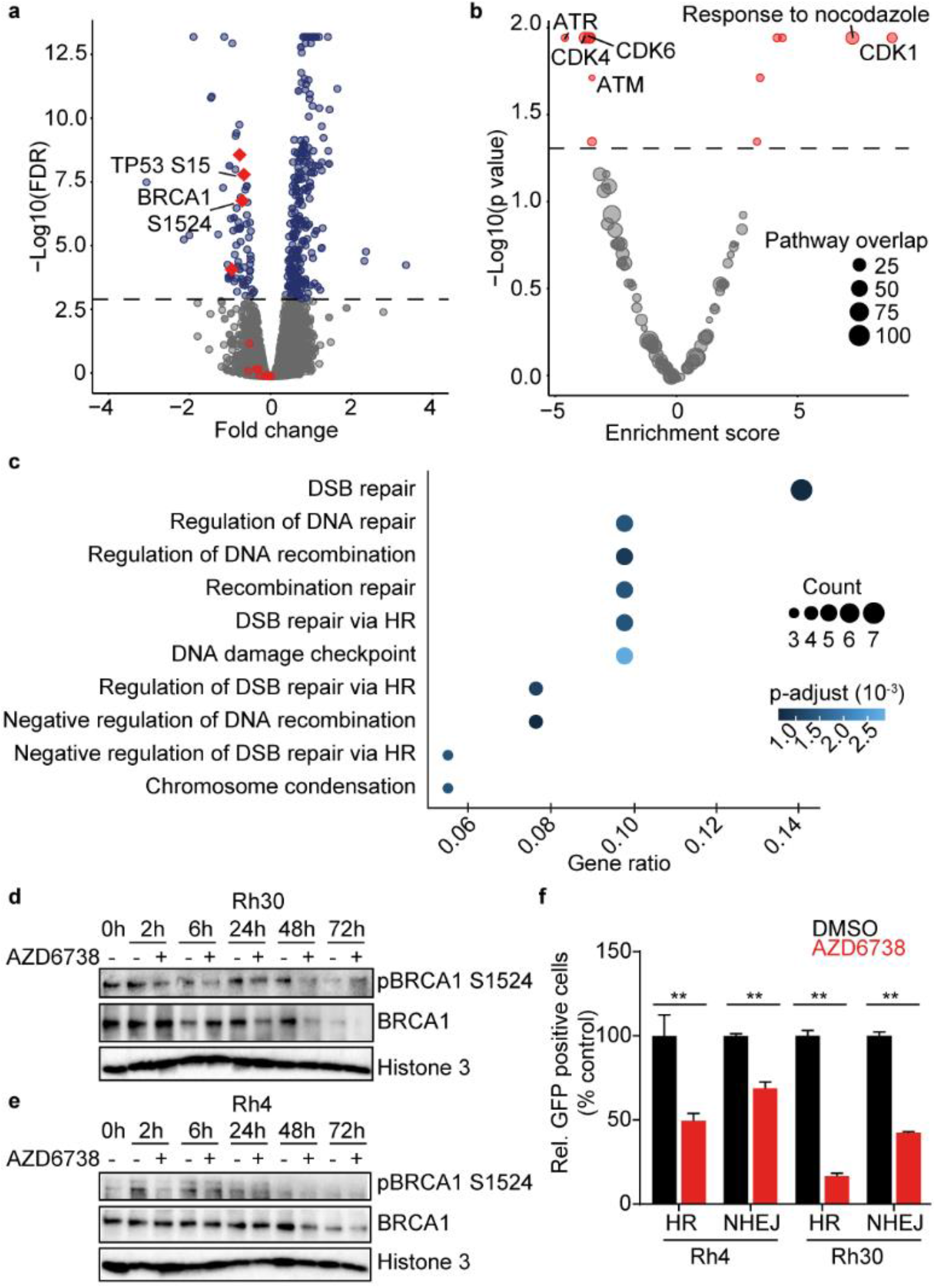
Pharmacological ATR inhibition leads to reduced BRCA1 activation and repressed homologous recombination in alveolar rhabdomyosarcoma cells. **(a)** Volcano plot showing relative changes in phospho-peptide abundance in PAX3-FOXO1- expressing Rh30 cells after 2 hours of incubation with AZD6738 (750 nM) measured using LC-MS/MS proteomics (red, known ATR targets; dotted line indicating a false discovery rate of 0.001). **(b)** Volcano plot showing relative enrichment of molecular pathways in which differential phospho-peptide abundance was observed in cells treated with AZD6738 (750 nM) compared to DMSO-treated cells (dotted line indicating a false discovery rate of 0.05). **(c)** Cellular processes significantly enriched in differentially abundant phospho-peptides. **(d-e)** Western immunoblotting of BRCA1 pSer1524 and total BRCA1 in Rh30 (d) and Rh4 (e) cells during the course of AZD6738 treatment (histone 3 serves as loading control). **(f)** Relative activity of HR and NHEJ in Rh4 and Rh30 cells after incubation with 750nM AZD6738 (From left to right, *P*=0.003, *P*=1.7×10^−4^, *P*=2.7×10^−6^ and *P*=2.1×10^−6^; two-sided student’s t-test; error bars represent standard error of the mean).

### ATR inhibition sensitizes cells to PARP inhibition and cisplatin in vitro

Based on the observation that ATR inhibition led to reduced BRCA1 activation and repressed HR pathway activity in rhabdomyosarcoma cells (Fig. 6), we hypothesized that the reduced replication stress-induced DNA damage repair via HR may increase dependence on PARP-mediated base excision repair (BER), as observed in *BRCA1*- deficient cancers^10^. Indeed, shRNA-mediated BRCA1 knock down in alveolar rhabdomyosarcoma cells with three independent shRNAs led to increased sensitivity to PARP inhibition, with IC_50_ for olaparib changing from 39.4 µM for shGFP-expressing cells to 5.01 µM, 7.19 µM and 15.84 µM for three shRNAs targeting BRCA1, respectively (Extended data Fig. 7a-b). This suggests that BER is required in rhabdomyosarcoma cells in the absence of BRCA1 activity. Consistently, BRCA1 knock down also sensitized rhabdomyosarcoma cells to ATR inhibition (Extended data Fig. 7c). We hypothesized that due to its effect on BRCA1 activity, pharmacological ATR inhibition could sensitize rhabdomyosarcoma cells to PARP inhibition as well as cisplatin, a cytotoxic agent also known to be more effective in the context of HR deficiency^10^. Indeed, significant synergy of combined AZD6738 and olaparib as well as combined AZD6738 and cisplatin treatment was detected by Excess over Bliss analysis (Fig. 7a-c and Extended data Fig. 7d-i). Thus, ATR inhibition sensitized rhabdomyosarcoma cells to olaparib and cisplatin, creating clinically exploitable opportunities for combination therapy.

**Fig. 7.**
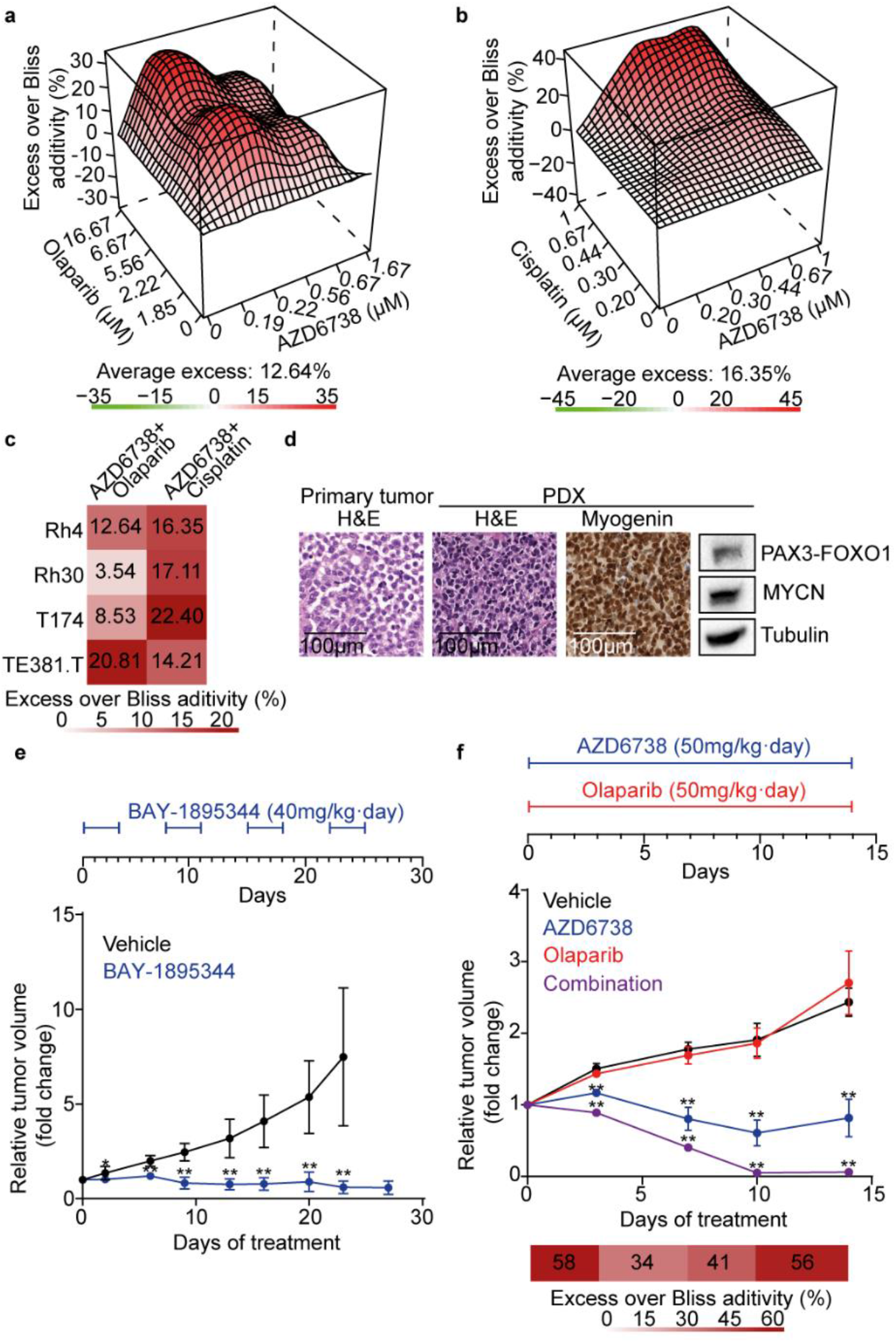
ATR inhibition sensitizes patient-derived alveolar rhabdomyosarcoma xenografts to PARP inhibition *in vivo*. **(a)** Excess over Bliss analysis of combined treatment with olaparib and AZD6738 in Rh4 cells (*n*=3). **(b)** Excess over Bliss analysis of combined treatment with cisplatin and AZD6738 in Rh4 cells. **(c)** Bliss score for four rhabdomyosarcoma cell lines treated with AZD6738 combined with olaparib or cisplatin. **(d)** Representative histological images and western immunoblot for PAX3-FOXO1 and MYCN for the alveolar rhabdomyosarcoma patient-derived xenograft (PDX) model used. **(e)** Tumor volume over time of mice harboring the alveolar rhabdomyosarcoma PDX model subcutaneously xenografted and treated with BAY-1895344 twice daily by oral gavage at 40mg/kg body weight per day on a 3 days on/4 days off schedule as compared to vehicle control treated mice (n=7 mice per group; top, timeline of the drug schedule). **(f)** Tumor volumes over time of mice harboring the alveolar rhabdomyosarcoma PDX model subcutaneously xenografted and treated with AZD6738, olaparib or both AZD6738 and olaparib compared to vehicle control treated mice (n=4 mice per group; bottom, excess over Bliss additivity for each treatment timepoint, \*\**P*<0.01; two-sided student’s t-test; error bars represent standard deviations).

### Combined inhibition of ATR and PARP has synergistic anti-tumor effects in patient-derived alveolar rhabdomyosarcoma models

We next sought to explore the effect of single agent ATR inhibition and its combination with olaparib in patient-derived PAX3-FOXO1-expressing alveolar rhabdomyosarcoma xenografts (PDX). Notably, we used a PAX3-FOXO1-expressing PDX model derived from a patient still undergoing treatment of a chemotherapy resistant alveolar rhabdomyosarcoma relapse at our hospital at the time the experiments outlined here were performed. Histological analysis of the PDX and matching patient tumor confirmed that the PDX model adequately reflected alveolar rhabdomyosarcoma histologically and expressed PAX3-FOXO1 and MYCN (Fig. 7d). In line with our observations *in vitro*, single-agent AZD6738 or BAY-1895344 treatment led to significant reductions in tumor burden over time in mice harboring the alveolar rhabdomyosarcoma PDX (Fig. 7e-f). Neither AZD6738 nor BAY-1895344 alone led to significant reduction in mouse weight (Extended data Fig. 8), indicating drug tolerability. Addition of olaparib to AZD6738 treatment significantly potentiated the anti-tumor effects, leading to full regression of the PDX (Fig. 7f). Loss of mouse weight after 10 days of combined AZD6738 and olaparib treatment, however, indicated increased toxicity compared to single agent treatment (Extended data Fig. 8b). Thus, combined pharmacological inhibition of ATR and PARP has synergistic activity against preclinical models of PAX3-FOXO1-expressing alveolar rhabdomyosarcomas, which may be clinically translatable.

### Compassionate use of ATR inhibitors in patients with PAX3-FOXO1-expressing rhabdomyosarcomas is clinically feasible and tolerable

Pediatric clinical trials with ATR inhibitors are currently being prepared. Encouraged by the promising anti-tumor activity of combined AZD6738 (ceralasertib) and olaparib treatment observed in the PDX model (Fig. 7f), which was derived from an 18 year old female patient undergoing treatment of a highly chemo-resistant PAX3-FOXO1- expressing alveolar rhabdomyosarcoma 5^th^ relapse at the Charité University Hospital in Berlin, we decided for a compassionate use of ceralasertib and olaparib in this patient. Prior to the compassionate use treatment, the patient had been treated according to the recommendations of the registry for soft tissue sarcoma and other soft tissue tumors in children, adolescents, and young adults (CWS-SoTiSaR). All conventional treatment options had been exhausted, while the patient’s tumor continued to progress during treatment. We initiated oral treatment with ceralasertib according to dose and schedule currently investigated in adult clinical trials (Extended data Fig. 9, NCT03682289, NCT03428607, NCT03462342 among others). Apart from supportive pain medication with non-opioid analgesics, no additional drugs were administered throughout the duration of treatment. The patient was closely clinically monitored on a bi-weekly basis, but standardized tumor imaging was not feasible due to the late stage of disease. Notably, the patient reported marked reduction of pain in tumor regions (back and abdomen) from 7 (severe pain) to 3 (mild pain) on a numeric rating scale of 10 after one week of combined ceralasertib and olaparib treatment. Apart from known on-target off-tumor toxicity of ceralasertib on blood counts, in particular thrombocytopenia grade 1 according to the Common Terminology Criteria for Adverse Events (CTCAE; Extended data Fig. 9), no considerable side effects were observed nor reported by the patient over the course of treatment. In parallel, we tested the anti-tumor efficacy of the ATR inhibitor BAY- 1895344 against primary patient-derived cells from another patient with relapsed PAX3- FOXO1-expressing alveolar rhabdomyosarcoma, hospitalized at the Memorial Sloan Kettering Cancer Center in New York (USA). Patient-derived cells were sensitive to inhibition of ATR by BAY-1895344 (Extended data Fig. 10a), leading us to begin the compassionate use of BAY-1895344 in this patient. The patient received 15 mg of BAY- 1895344 *per os* twice a day for 3 days followed by 4 days of drug pause for 3 weeks. After demonstrating tolerability, doses were increased to 18 mg twice a day (Extended data Fig. 10b). The patient refused further assessment using imaging, but reported clinically relevant improvement in pain and laboratory signs that may reflect response, such as decreased lactate dehydrogenase (LDH) levels (Extended data Fig. 10b). Even though both patients progressed within one month after ATR inhibitor treatment, the two compassionate use cases presented here provide promising evidence that ATR inhibitors ceralasertib and BAY-1895344 may present clinically tolerable and feasible treatment options for patients suffering from alveolar rhabdomyosarcomas.

## Discussion

We have found that PAX3-FOXO1, a pathognomonic fusion oncogene in alveolar rhabdomyosarcoma, confers a synthetic lethal dependency on ATR-mediated DNA damage repair. Consistent with previous reports of other oncogenic fusion genes inducing replication stress in Ewing sarcoma^33^, expression of PAX3-FOXO1 was sufficient to increase replication stress, which required both DNA damage repair and DNA damage signaling, resulting in apoptosis if impaired by the selective inhibition of ATR, or ATR- pathway members. Both untransformed mouse myoblast cells and rhabdomyosarcoma cells engineered to express PAX3-FOXO1, as well as PAX3-FOXO1-expressing rhabdomyosarcoma cell lines, accumulated unrepaired DNA damage and underwent apoptosis upon treatment with selective inhibitors of ATR signaling. These effects, observed specifically in PAX3-FOXO1-expressing rhabdomyosarcoma cells, were associated with decreased phosphorylation of BRCA1 and homologous recombination activity, and were accompanied by induction of genomic instability, increased G2/M accumulation and apoptosis. In turn, single-agent treatment with structurally diverse inhibitors of ATR exhibited potent antitumor activity against high-risk patient-derived alveolar rhabdomyosarcoma models. Moreover, decreased BRCA1 and homologous recombination activity through pharmacological ATR inhibition sensitized cells to PARP inhibition. When combined, ATR and PARP inhibitors exhibited strong antitumor activity against patient-derived rhabdomyosarcoma models. Lastly, compassionate use of ATR inhibitors in two cases of relapsed, PAX3-FOXO1-driven, alveolar rhabdomyosarcoma indicated clinical feasibility and tolerability.

Human cancers require active DNA damage repair for survival. As a result, selective inhibitors of ATR-mediated DNA damage repair signaling have been used to target tumors with intrinsic deficiencies in DNA repair or high abundance of DNA damage. Dissecting the molecular mechanisms of susceptibility to ATR inhibitors has been the subject of extensive investigations in the past years^46^. We and others have found inducers of ATR inhibitor susceptibility, such as PGBD5 recombinase activity in embryonal tumors^21^, oncogene-induced replication stress, ATM loss, and TP53 deficiency^20,47^. Our current work revealed a specific synthetic lethal dependency conferred by high steady-state replication stress in alveolar, PAX3-FOXO1-expressing rhabdomyosarcoma. Pharmacological inhibition of DNA damage signaling kinases exhibited a specific response profile, with ATR- and CHK1-selective inhibitors showing enhanced replication stress-dependent anti-tumor activity. Notably, CHK1, a downstream target of ATR, is inhibited by prexasertib, which is currently being clinically investigated in combination with chemotherapy for patients with relapsed rhabdomyosarcoma (NCT04095221). Given their varied potency and selectivity, it is possible that other selective DNA damage signaling inhibitors can also effectively target replication stress-induced synthetic lethal dependencies in rhabdomyosarcoma. In line with replication stress-dependent anti-tumor activity of ATR inhibitors, our genome-wide CRISPR-based screen identified potent replication stress inducers such as *MYC* as sensitizers to ATR inhibition. Because ATR is also activated by specific DNA structures such DNA-RNA hybrid R-loops, which can be the cause of oncogene-induced DNA replication stress^33^, the apparent selective activity of ATR inhibitors in PAX3-FOXO1-expressing cells may also be due to the formation of such structures. We provide evidence that PAX3-FOXO1 expression, at least in part, contributes to replication stress and hypersensitivity to ATR inhibition, which is consistent with previous reports of fusion oncogene-induced replication stress in Ewing sarcoma^32,33^. High *MYCN* expression was also detected in PAX3-FOXO1-expressing cells and was positively associated with ATR inhibitor sensitivity. *MYCN* has been described as a direct target of PAX3-FOXO1 and is itself a potent inducer of replication stress^48^. Thus, MYCN likely also contributes to high replication stress and hypersensitivity to ATR inhibition in PAX3-FOXO1-expressing cells. Alternative lengthening of telomeres (ALT) is also frequently observed in embryonal rhabdomyosarcoma not expressing PAX3-FOXO1^49-51^. Due to the lack of models for robust induction of ALT in human cells, we currently cannot assess the contribution of ALT to ATR inhibitor sensitivity in rhabdomyosarcoma. Nevertheless, the susceptibility of PAX3-FOXO1-expressing tumors to selective ATR inhibitors is expected to cooperate with oncogene-, ALT-associated and other sources of replication stress.

ATR is essential for intra-S phase and G2/M checkpoint activation^13,16,25,52^. When checkpoints are constitutively active, cells can undergo checkpoint adaptation to continue proliferating despite the presence of DNA damage^53,54^. We anticipate that susceptibility to ATR inhibitors may also depend on tumor-specific mechanisms of checkpoint adaptation. While little is known about checkpoint adaptation in human cells, CHK1 and CDK1 can promote checkpoint adaptation by mediating forced mitotic entry^55,56^. Inhibition of ATR could exacerbate the effect of checkpoint adaptation by suppressing checkpoint activation. Consistently, we observed accumulation of cells in G2/M and increased activation of CDK1 targets in our phosphoproteomic profiling after ATR inhibition. In line with checkpoint adaptation promoting DNA damage accumulation and genomic instability^54^, we observed high degrees of genomic instability in PAX3-FOXO1- expressing cells treated with ATR inhibitors. Intriguingly, PAX3-FOXO1 can itself promote checkpoint adaptation in rhabdomyosarcoma cells through induction of *PLK1* expression, which in turn activates CDK1 and forced mitotic entry^57^. It is tempting to speculate that ATR dependency in PAX3-FOXO1-expressing rhabdomyosarcoma could also be due to checkpoint adaptation.

Even though results of clinical trials with ATR inhibitors in adults have shown promising single agent antitumor activity in various tumor entities, many patients progress or relapse after some time^58,59^. Thus, identifying molecular mechanisms of ATR inhibitor resistance is of paramount clinical importance, as it may enable the identification of clinical biomarkers that help predict ATR inhibitor susceptibility and can be used to monitor resistance development. Our genome-wide CRISPRa screen in rhabdomyosarcoma cells identified the *FOS* family of transcription factors as repressors of replication stress, leading to decreased sensitivity towards ATR inhibition in rhabdomyosarcoma cells. Apart from their role in DNA damage repair regulation^35,36^, the *FOS* gene family are known negative regulators of myoblast differentiation and can directly repress *MYOD, MYOG* and *PAX7* expression both in primary myoblasts and rhabdomyosarcoma cell^60,61^, suggesting that rhabdomyosarcoma cell differentiation may represent an additional mechanism through which *FOS* gene family members reduce sensitivity to ATR inhibition. It remains to be determined, however, how the intrinsic ability to differentiate or the pre-existence of cells in different differentiation states contributes to ATR inhibitor resistance. It is well described that transitions between differentiation states in cancers occurs^62^, but the underlying mechanisms leading to epigenetic and transcriptional reprogramming are still poorly understood. Based on our observations, we anticipate that studying trans-differentiation between stable or unstable epigenetic cellular states in rhabdomyosarcoma may improve our understanding of rhabdomyosarcoma resistance to therapy.

In conclusion, we here present preclinical evidence and clinical observations supporting a molecularly targeted therapeutic option for an aggressive subset of rhabdomyosarcomas, for which current treatment options have been exhausted and prognosis remains dismal. Our findings warrant the future investigation of ATR inhibitors in clinical trials for children with PAX3-FOXO1-expressing alveolar rhabdomyosarcoma, the majority of which should be characterized by ATR-signaling dependencies.

## Materials and Methods

### Study design

The purpose of this study was to examine the effects of ATR inhibition in preclinical models of rhabdomyosarcoma and identify potential biomarkers to select patients that could benefit from small molecule ATR inhibitor treatment. We first determined the inhibitory activity of the ATR inhibitors in rhabdomyosarcoma cell models, and compared these cells based on known determinants of ATR inhibition sensitivity, as well as PAX3-FOXO1, a molecular feature of alveolar rhabdomyosarcomas. We analyzed the effects of AZD6738 treatment on genomic instability (including double strand break formation, micronucleation, and apoptosis) and on protein phosphorylation. All *in vitro* experiments were performed in at least three technical replicates in at least two cell models of each meolecular rhabdomyosarcoma subgroup (PAX3-FOXO1-expressing vs. embryonal rhabdomyosarcoma). Outliers were not excluded unless technical errors were present. For the CRISPRa screen, we used only one cell line and at least three independent sgRNAs per gene. All sgRNAs of interest were validated in independent experiments. For the analysis of phosphoproteomic changes after ATR inhibition, we used three independently grown biological replicates of the same rhabdomyosarcoma cell line. For *in vivo* testing, sample size was decided based on previous experience with the models. Animals euthanized before the end of the experiment, due to excessive tumor growth or loss of body weight, were included in the analysis. For patient use of the drugs, we consulted with the company the appropriate dose regimen based on current clinical trials. The researchers and patients were not blinded during the experiments.

### Reagents

All reagents were obtained from Carl Roth (Karlsruhe, Germany) unless otherwise indicated. Oligonucleotide primers were obtained from Eurofins Genomics (Luxemburg, Luxemburg, Table 1). A list of antibodies and their catalog numbers can be found in Table 2. AZD6738 (celarasertib) was provided by Astra Zeneca (Cambridge, UK). All drugs were dissolved in Dimethylsulfoxid (DMSO) and stored at 10 mM concentrations at -20 °C.

### Plasmid constructs

Human *PAX3-FOXO1* cDNA was PCR-amplified and isolated from a plasmid gifted by Prof. Beat Schäfer. *PAX3-FOXO1* cDNA was cloned into pENTR1A (Thermo Fisher) using the restriction enzymes SalI and NotI (New England Biolabs), and cloned into a pInducer20 (Addgene #44012) using the Gateway strategy and the manufacturer’s protocol (Thermo Fisher). pLKO.1 shRNA plasmids targeting BRCA1 (TRCN0000009823, TRCN0000010305, TRCN0000039834), and control targeting GFP (shGFP) were obtained from the RNAi Consortium (Broad Institute). Plasmid containing an inducible shRNAs targeting PAX3-FOXO1 (cloned in the pRSI backbone) were a kind gift from Prof. Beat Schäfer.

### Cell culture

Rh41, TE441.T, Kym1 and Rh18 cells were a kind gift from Prof. Simone Fulda. The remaining human tumor cell lines were obtained from the American Type Culture Collection (ATCC, Manassas, Virginia). The absence of *Mycoplasma sp*. contamination was determined using a Lonza (Basel, Switzerland) MycoAlert system. Rh4, Rh30, Rh41, RD, T174, TE381.T, TE441.T, C2C12, HEK293T, RPE1-hTERT and BJ cell lines were cultured in Dulbecco’s Modified Eagle’s Medium (DMEM, Thermo Fisher, Waltham, Massachusetts, USA) supplemented with 10% fetal calf serum (Thermo Fisher) and penicillin/streptomycin (Thermo Fisher). Rh18 and Kym1 cells were cultured in Roswell Park Memorial Institute (RPMI)-1640 (Thermo Fisher) supplemented with 10% fetal calf serum and penicillin/streptomycin. Twice per week, cells were washed with phosphate-buffered saline (PBS), incubated in trypsin (Thermo Fisher) for five minutes sedimented at 500 g for 5 minutes and a fraction was cultured in fresh media. Cells were kept in culture for a maximum of 30 passages. Resuspended cells were counted by mixing 1:1 with 0.02 % trypan blue in a BioRad (Hercules, CA, USA) TC20 cell counter.

### Lentiviral transduction

Lentivirus were produced as previously described^63^. In short, HEK293T cells were transfected using TransIT-LT1 (Mirus, Madison, Wisconsin, USA) in a 2:1:1 ratio of lentiviral plasmid, psPAX2 and pMD2.G plasmids following the TransIT-LT1 manufacturer’s protocol. Viral supernatant was collected 48h and 72h after transfection, pooled, filtered and stored at -80 °C. Cells were transduced for one day in the presence of 8µg/mL polybrene (Sigma Aldrich).

### CRISPRa screening and sequencing

The genome-wide CRISPRa screen was performed as described in Konermann *et al*.^34^. Briefly, Rh4 cells were transduced with the lentiMPH v2 plasmid (Addgene #89308) and selected with hygromycin for 10 days (Thermo Fisher). Next, cells were transduced with the sgRNA library at a multiplicity of infection (MOI) of < 0.3, ensuring at least 500 cells to be transduced with each sgRNA-encoding plasmid on average. After selection with blasticidin (Thermo Fisher) for 7 days, cells were separated in two groups, one group was incubated in the presence of AZD6738 at 750 nM concentration and the other group was incubated in the presence of DMSO. Genomic DNA was extracted and the sgRNA amplified using PCR and barcoded for Illumina sequencing. Sequencing was performed on a NextSeq500 with Mid Output, with a read length of 1 x 81bp +8 bp Index and 20% PhiX Control v3. Samples were demultiplexed using flexbar^64^ and analyzed using MaGECK^65^.

### Dependency analysis using DepMap

To study dependency to DNA damage proteins, we used the dataset CRISPR Avana 20Q2 publicly available in the online resource DepMap^27^. Cell lines were grouped by tissue of origin, and groups with only one cell line were excluded from the analysis. For rhabdomyosarcoma, we subdivided the group into subgroups in order to differentiate alveolar, embryonal and other rhabdomyosarcomas.

### Cell viability

Cell viability was assessed using CellTiter-Glo (Promega, Madison, Wisconsin, USA). Briefly, for CellTiter-Glo measurement, 1,000 cells were seeded in white, flat-bottom, 96-well plates (Corning, Corning, NY, USA). After 24 hours, drugs were added to the medium and cells were incubated for 72 hours. CellTiter-Glo luminescent reagent was added according to the manufacturers protocol, and the luminescence signal measured on a Glowmax-Multi Detection System (Promega). To evaluate if a combination of drugs is synergistic, cells were simultaneously treated with varying concentrations of drugs, and cell viability was measured with CellTiter-Glo. Synergism scores were obtained using the R package SynergyFinder^66^. For patient-derived cells, dose-response studies were performed using 11 doubling dilutions in triplicate with 100µM, 10µM and 1µM as the highest concentration (5uL in 10% DMSO). After cell seeding (45uL), the plates were placed in a controlled environment at 37°C and 5% CO2 for 3 days. Then, 5µL Alamar Blue (AB) were added using the Multidrop™ Combi. They were further incubated for 1 day, and the fluorescence intensity was read on a Cytation 5 Cell Imaging Multi-Mode Reader (Biotek).

### Immunoblotting

Whole-cell protein lysates were prepared by lysing cells in Radioimmunoprecipitation assay buffer (RIPA) supplemented with cOmplete Protease inhibitor (Roche, Basel, Switzerland) and PhosphStop (Roche). To enrich in nuclear proteins, samples were prelysed in a soft lysis buffer (5mM PIPES ph=8.0, 85mM KCl, 0.5%NP-40), followed by the lysis with RIPA buffer. Protein concentrations were determined by bicinchoninic acid assay (BCA, Thermo Fisher). 10 µg of protein were denatured in Laemmli buffer at 95 °C for 5 minutes. Lysates were loaded onto 16%, 10% Tris-Glycin (Thermo Fisher) or 3-8% Tris-Acetate gels (Thermo Fisher) for gel electrophoresis depending on the protein sizes of interest. Proteins were transferred onto Polyvinylidenfluorid (PVDF) membranes (Roche), blocked with 5% dry milk for 1 hour and incubated with primary antibodies overnight at 4°C, then secondary antibodies for 1 hour at room temperature. Chemiluminescent signal was detected using Enhanced chemiluminescence (ECL) Western Blotting Substrate (Thermo Fisher) and a Fusion FX7 imaging system (Vilber Lourmat, Marne-la-Vallée, France). Quantification was performed with ImageJ.

### Immunofluorescence

Cells were grown at the desired confluency on a glass coverslide for 24h (micronuclei quantification) and treated with 1000ng/mL doxycycline for another 48h (for the corresponding experiment). Cells were washed with PBS three times and fixed for 10 minutes with 4% paraformaldehyde, washed with PBS three times and permeabilized with PBS containing 0.1% Triton-X100. For micronuclei detection, cells were mounted on a slide with DAPI-containing mounting media (Vectashield, Vec-H-1000). For immunofluorescence, cells were blocked for 40 minutes with 5% BSA in PBS, incubated overnight at 4^a^C with the primary antibody (yH2AX; Millopore (#05-636), washed three times with PBS-T (0.05% Tween-20 in PBS), incubated for 1 hour in the dark at room temperature with the secondary antibody (Dianova, 715-096-150), washed three times with PBS-T and mounted on a slide with DAPI-containing mounting media. Cells were imaged using an ECHO Revolve microscope and quantified using ImageJ.

### RT-qPCR

RNA from cell lines was extracted using RNeasy mini kit (QIAGEN). Synthesis of cDNA was performed using Transcription First Strand cDNA Synthesis kit (Roche). 50 ng of cDNA were combined with the corresponding primers (Table 1), and SG qPCR Master Mix (Roboklon, Berlin, Germany), keeping the mixture and cycling conditions recommended by the manufacturer.

### Fluorescence-activated cell sorting (FACS)

For cell cycle analysis, cells were incubated with 5-Ethynyl-2′-deoxyuridine (EdU) for 2 hours and fluorescent labeling was performed with the Click-IT EdU Alexa Fluor 488 Flow Cytometry Assay kit (Thermo Fisher), according to the manufacturer’s description. Terminal deoxynucleotidyl transferase dUTP nick end labeling (TUNEL) was performed using the APO-BrdU TUNEL Assay Kit (Thermo Fisher), according to the manufacturer’s descriptions. Cell death was assessed by measuring caspase 3 cleavage using a CellEvent Caspase3/7 Green Flow Cytometry kit (Thermo Fisher), according to the manufacturer’s descriptions. Stained cells were measured on a BD LSR Fortessa flow cytometer (BD Biosciences, Franklin Lakes, NJ, USA).

### DNA damage repair activity assay

All the plasmids were obtained from Addgene (pDRGFP #26475; pimEJ5GFP #44026 ; pCBA-SceI #26477; pCAG-FALSE #89689;pCAGGS-mCherry #41583). The protocol was adapted from the plasmid depositors’^67,68^. Briefly, cells were co-transfected with pCBA-SceI and pDRGFP, or pCBA-SceI and pimEJ5GFP to analyze homologous recombination and non-homologous end joining activity, respectively. As a negative control, pCBA-SceI was substituted with the empty backbone pCAG-FALSE. Transfection efficiency was calculated using cells transfected with pCAGGS-mCherry. Two days after transfection, cells were trypsinized, washed twice with PBS and fluorescence measured with flow cytometry.

### Phosphoproteomics sample preparation

Rh30 cells were cultured for two weeks in the presence of stable isotope labeling with amino acids (SILAC) media in DMEM, 10% dialysed fetal calf serum, 1% Proline, 1% Glutamine, 0.025% ^8^Lysine and ^10^Arginine (“Heavy”) or ^0^Lysine and ^0^Arginine (“Light”). After labelling, cells were incubated in the presence of AZD6738 750 nM or DMSO for two hours in biological triplicates. Cells were harvested, resuspended and combined in 400 µL of 8 M urea and 0.1 M Tris-HCl, pH 8. Proteins were reduced in 10 mM dithiothreitol (DTT) at room temperature for 30 minutes and alkylated with 50 mM iodoacetamide (IAA) at room temperature for 30 minutes in the dark. Proteins were first digested by lysyl endopeptidase (LysC) (Wako Pure Chemical Industries, Ltd., Osaka, Japan) at a protein-to-LysC ratio of 100:1 (w/w) at room temperature for 3 hours. Then, the sample solution was diluted to final concentration of 2 M urea with 50 mM ammonium bicarbonate (ABC). Trypsin (Promega) digestion was performed at a protein-to-trypsin ratio of 100:1 (w/w) under constant agitation at room temperature for 16 hours. Tryptic digests corresponding to 200 µg protein per condition were desalted with big C18 Stage Tips packed with 10 mg of ReproSil-Pur 120 C18-AQ 5 µm resin (Dr. Maisch GmbH, Ammerbuch, Germany). Peptides were eluted with 200 µL of loading buffer (80% ACN (v/v) and 6% TFA (v/v). Phosphopeptides were enriched using a microcolumn tip packed with 0.5 mg of TiO2 (Titansphere, GL Sciences, Tokyo, Japan)^69^. The TiO2 tips were equilibrated with 20 μL of the loading buffer via centrifugation at 100 g. 50 μL of the sample solution was loaded on a TiO2 tip via centrifugation at 100 g and this step was repeated until the sample solution was loaded. The TiO2 column was washed with 20 μL of the loading buffer, followed by 20 μL of washing buffer (50% ACN (v/v) and 0.1% TFA (v/v)). The bound phosphopeptides were eluted using successive elution with 30 μL of elution buffer 1 (5% ammonia solution), followed by 30 μL of elution buffer 2 (5% piperidine)^70^. Each fraction was collected into a fresh tube containing 30 μL of 20% formic acid. 3 μL of 100% formic acid was added to further acidify the samples. The phosphopeptides were desalted with C18 Stage Tips prior to nanoLC-MS/MS analysis.

### NanoLC-MS/MS analysis

Peptides were separated on a 2 m monolithic column (MonoCap C18 High Resolution 2000 (GL Sciences), 100 µm internal diameter x 2,000 mm at a flow rate of 300 nL/min with a 5 to 95 % acetonitrile gradient on an EASY-nLC II system (Thermo Fisher Scientific). 240-min gradient was performed for phosphoproteome analyses. A Q Exactive plus instrument (Thermo Fisher Scientific) was operated in the data dependent mode with a full scan in the Orbitrap followed by top 10 MS/MS scans using higher-energy collision dissociation (HCD). For whole proteome analyses, the full scans were performed with a resolution of 70,000, a target value of 3×10^6^ ions and a maximum injection time of 20 ms. The MS/MS scans were performed with a 17,500 resolution, a 1×10^6^ target value and a 20 ms maximum injection time. For phosphoproteome analyses, the full scans were performed with a resolution of 70,000, a target value of 3×10^6^ ions and a maximum injection time of 120 ms. The MS/MS scans were performed with a 35,000 resolution, a 5×10^5^ target value and a 160 ms maximum injection time. Isolation window was set to 2 and normalized collision energy was 26.

Raw data were analyzed and processed using MaxQuant (v1.5.1.2)^71^. Search parameters included two missed cleavage sites, fixed cysteine carbamidomethyl modification, and variable modifications including L-[^13^C_6_,^15^N_4_]-arginine, L-[^13^C_6_,^15^N_2_]-lysine, methionine oxidation, N-terminal protein acetylation, and asparagine/glutamine deamidation. In addition, phosphorylation of serine, threonine, and tyrosine were searched as variable modifications for phosphoproteome analysis. The peptide mass tolerance was 6 ppm for MS scans and 20 ppm for MS/MS scans. Database search was performed using Andromeda^72^ against uniprot-human 2014-10 with common contaminants. False discovery rate (FDR) was set to 1% at both peptide spectrum match (PSM) and protein level. The ‘re-quantify’ and ‘match between runs’ functions were enabled. Phosphorylation sites were ranked according to their phosphorylation localization probabilities (P) as class I (P > 0.75), class II (0.75 > P > 0.5) and class III sites (P < 0.5), and only class I sites were used for further analyses. Data normalization was performed using the default settings of the R package DEP^73^. In short, peptides not identified in at least two replicates in both conditions were removed. Intensity values were normalized based on the variance stabilizing transformation, and missing values were imputed using random draws from a Gaussian distribution centered around a minimal value (q= 0.01). For pathway enrichment analysis, we used a single sample gene set enrichment analysis (ssGSEA) as previously described^38^, ranking genes according to their fold change. For gene ontology (GO) analysis, we followed the ClusterProfiler R package^74^. P-values were calculated using hypergeometric distribution (one-sided Fisher exact test) and corrected for multiple comparisons (Holm-Bonferroni method), selecting phosphopeptides with a fold change>1 or <-1 and a FDR<0.01, and reporting the top 10 GO terms enriched in the subset.

### Patient-derived xenograft (PDX) treatment

The establishment of PDX models was conducted as previously described^75^ in collaboration with Experimental Pharmacology & Oncology GmbH (EPO, Berlin, Germany). All experiments were conducted according to the institutional animal protocols and the national laws and regulations. Tumor fragments from rhabdomyosarcoma patients were transplanted into NOD.Cg-Prkdcscid Il2rgtm1Sug/JicTac mice (Taconic, Rensselaer, NY, USA). Tumor growth was monitored with caliper measurements. Tumor volume was calculated with the formula length x width^2 / 2. PDX were serially transplanted in mice at least three times prior to the experiments. Mice were randomized into four groups with at least 3 mice to receive AZD6738 (50 mg/kg day, oral), cisplatin (2mg/kg twice weekly), olaparib (50 mg/kg day, oral), a combination of AZD6738 and either cisplatin or olaparib, or vehicle. For in vivo treatment, AZD6738 was dissolved in DMSO at 62.5 mg/ml and mixed 1:10 in 40% propylene glycol and 50% sterile water, resulting in a final AZD6738 concentration of 6.25 mg/ml. Cisplatin was dissolved in phosphate-buffered saline and administered intraperitoneally. Olaparib was dissolved in 4% DMSO, 30% polyethylene glycol 300 and sterile water. For the BAY-1895344 study, mice were administered 40mg/kg body weight on a 3 days on/ 4 days off regime twice daily (orally). BAY-1895344 was dissolved in 60% polyethylene glycol 400, 10% ethanol and 30% water to a 4mg/ml solution. Mice were sacrificed by cervical dislocation once the tumor volume exceeded 2,000 mm^3^ or body weight loss was higher than 10%. Solutions in which the drugs were dissolved were used as vehicle controls respectively.

### Compassionate AZD6738 and BAY-1895344 use

Oral treatment with AZD6738, olaparib and BAY-1895344 were provided according to a compassionate-use protocol. Drugs were provided by AstraZeneca and Bayer, respectively. Treatment regimens were based on regimens currently tested in clinical trials, and the patients/guardians provided written informed consent to participate in the compassionate use protocols and this case report. There was no commercial support for this study. Tumor material was collected according to the clinical protocol. Peripheral blood samples were collected in vacutainer tubes ± Ethylenediaminetetraacetic acid (EDTA) or ± Heparin. Simultaneous PET/MRI was performed with a 3T MRI/PET Magnetom Biograph mMR hybrid system (Siemens Healthcare, Erlangen, Germany; software vB20P), featuring avalanche photodiode and total imaging matrix coil technology. MR parameters included: MQ- Gradients: 45 mT/m maximum gradient amplitude; 200 T/m/s maximum gradient slew rate; LSO crystal; 4.3 mm transverse spatial resolution at FWHM at 1 cm; 15.0 kcps/MBq sensitivity at center; 13.8 kcps/MBq at 10 cm off-center. [^18^F]FDG was administered intravenously using a weight-adapted activity based on recommendations provided by the European Association of Nuclear Medicine (EANM). A test of blood glucose level was mandatory to assure that blood glucose level was ≤8.3 mmol/l. The PET scan was performed after an uptake time of ≥55 minutes in supine position from vertex to toes (3D mode; bed overlap, 53.3%). Among others, acquired MRI-sequences included axial T1 volumetric interpolated breath-hold examination (VIBE) using two-point Dixon fat-water separation (Dixon-VIBE) images following bodyweight-adapted administration of gadolinium-based MRI contrast agent (Gadovist^®^, Bayer, Leverkusen, Germany). Within the PET/MRI data generated, all areas with pathologically increased tracer uptake were identified and volumetrically assessed using the corresponding MRI-sequence. The MRI sequences were acquired on a MAGNETOM Skyra (Siemens) after application of 6.9 ml gadolinium-based MRI contrast agent (Bayer, Leverkusen, Germany). Tumor foci were measured in axial T1 volumetric interpolated breath-hold examination (VIBE) using two-point Dixon fat-water separation (Dixon-VIBE) images. Ultrasonography was performed with a GE logiq e9 R6 instrument (General Eclectics, Boston, USA).

### Statistical analysis

All statistical tests were done using GraphPrism7 (student’s two-sided t-test) or were part of the R package used for the analysis (MAGeCK, DEP, CePa).

## Extended data

**Extended Data Fig. 1**. Alveolar rhabdomyosarcoma cells express ATR and depend on expression of ATR/CHK1 pathway members.

**Extended Data Fig. 2**. PAX3-FOXO1-expressing alveolar rhabdomyosarcoma cells are hypersensitive to pharmacological ATR pathway inhibition.

**Extended Data Fig. 3**. Pharmacological ATR inhibition exacerbates replication stress and leads to genomic instability, apoptosis and cell cycle disruption.

**Extended Data Fig. 4**. Molecular factors associated with ATR inhibitor sensitivity in rhabdomyosarcoma cells.

**Extended Data Fig. 5**. Ectopic expression of PAX3-FOXO1 increases sensitivity to ATR inhibition in rhabdomyosarcoma cells.

**Extended Data Fig. 6**. FOSB, FOSL1 and FOSL2 expression induces resistance to ATR inhibition in rhabdomyosarcoma cells.

**Extended Data Fig. 7**. ATR inhibition synergizes with cisplatin and olaparib in rhabdomyosarcoma cell lines.

**Extended Data Fig. 8**. Combined treatment of mice with AZD6738 and olaparib has no remarkable toxicity in mice harboring alveolar rhabdomyosarcoma PDX models.

**Extended Data Fig. 9**. Compassionate use of AZD6738 and olaparib in a patient suffering from relapsed metastasized PAX3-FOXO1-expressing alveolar rhabdomyosarcomas.

**Extended Data Fig. 10**. Compassionate use of BAY-1895344 in a patient suffering from relapsed metastasized PAX3-FOXO1-expressing alveolar rhabdomyosarcomas.

**Table 1**. Oligonucleotide primers.

**Table 2**. Antibodies.

## Acknowledgments

We thank Experimental Pharmacology & Oncology GmbH for technical support. We thank Astra Zeneca for providing AZD6738 (ceralasertib) and olaparib for the compassionate use, as well as Emma Dean, Alan Lau, Andrew Pierce and Bienvenú Loembé for the fruitful discussions. We thank Bayer for providing BAY-1895344 and support for preclinical studies using that drug.

## Funding

The project that gave rise to these results received the support of a fellowship from “la Caixa” Foundation (ID 100010434). The fellowship code is LCF/BQ/EU18/11650037. A.G.H. is supported by the *Deutsche Forschungsgemeinschaft* (DFG, German Research Foundation) – 398299703 and the *Wilhelm Sander Stiftung*. This research was financially supported by the Charité 3R, Charité - Universitätsmedizin Berlin. A.G.H. is supported by the German Cancer Consortium (DKTK). A.G.H. is a participant in the BIH-Charité Clinical Scientist Program funded by the Charité – *Universitätsmedizin Berlin* and the Berlin Institute of Health. M.V.O. is supported by Cannonball Kids’ cancer and the NIH/NCI via K12 CA184746 and P30 CA008748. This work was supported by the *Deutsche Krebshilfe* (German Cancer Aid) – 70113870 and 70113871. We thank the patients and their parents for granting access to the tumor specimen and clinical information that were analyzed in this study.

## Author Contributions

H.D.G. and A.G.H. contributed to the study design, collection and interpretation of the data and wrote the manuscript. H.D.G., Y.B., J.v.S., G.I., K.I., N.T., R.C.G., I.M., F.P., C.Y.C., J.S., C.F., and B.L. performed experiments, analyzed data and reviewed this manuscript. H.D.G., K.He. and K.Ha. performed the analysis of the CRISPR screening. V.B., D.G., and J.R. collected and prepared PDX samples. A.W., A.E., M.Sc., G.S., P.H., M.K., P.M., M.Se., A.L., J.H.S., and M.O. contributed to study design. A.G.H. led the study design, to which all authors contributed.

## Competing interests

A.M.W. is employed by Bayer AG. A.G.H. has received research funding from Bayer AG.

## Data availability

All data is available upon request.

## Materials & Correspondence

Correspondence and requests for materials should be addressed to.

## Notes

### Competing Interest Statement

A.M. W. is employed by the Bayer AG.
A.G.H. has received research funding from Bayer AG.

