## Extended data for "Exploiting a PAX3-FOXO1-induced synthetic lethal ATR dependency for rhabdomyosarcoma therapy"

### PAX3-FOXO1-expressing alveolar rhabdomyosarcomas are hypersensitive to ATR inhibition

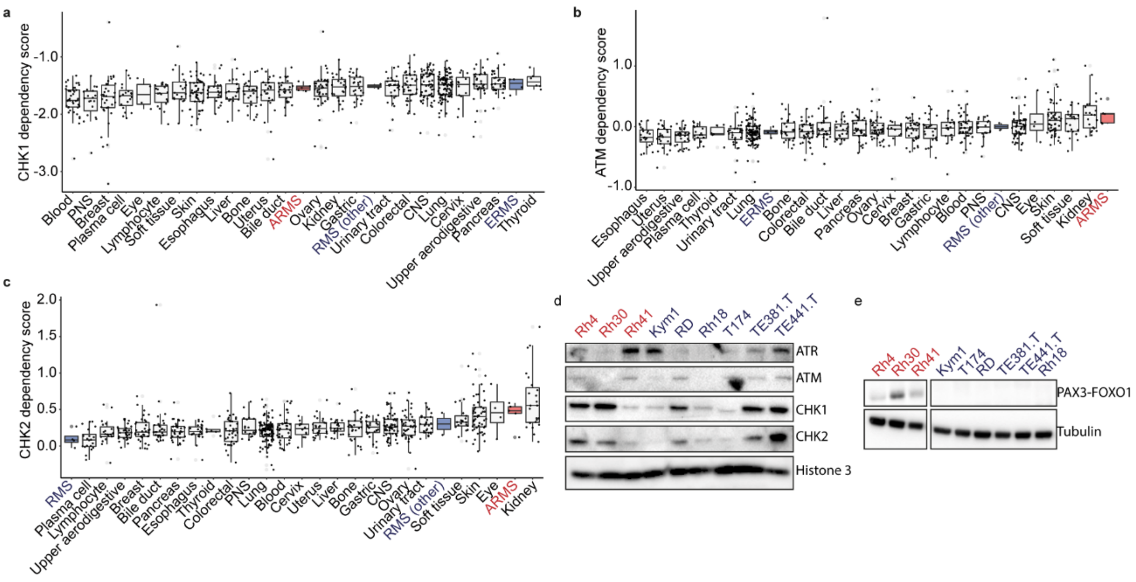


**Extended Data Fig. 1. Alveolar rhabdomyosarcoma cells express ATR and depend on expression of ATR/CHK1 pathway members.** **(a-c)** Dependency scores of several cancer types obtained from CRISPR (Avana) 20Q2 for CHK1 (a), ATM (b) and CHK2 (c). **(d)** Western immunoblot of ATR, ATM, CHK1 and CHK2 in rhabdomyosarcoma cell lines (Histone 3 as a loading control). **(e)** Western immunoblot of PAX3-FOXO1 in rhabdomyosarcoma cell lines. (Tubulin as a loading control). In red, alveolar rhabdomyosarcoma; in blue, other rhabdomyosarcomas.

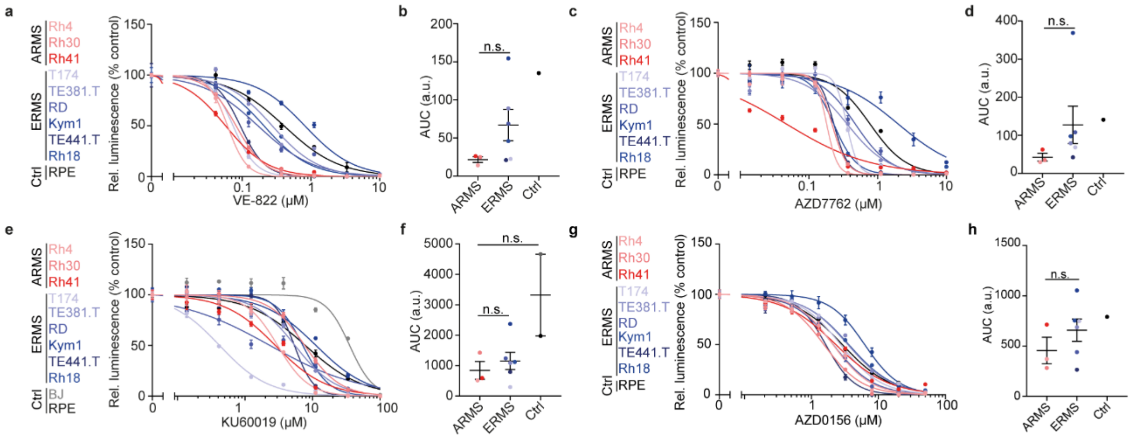


i

| Cell line | IC50 AZD6738 (µM) | IC50 BAY-1895344 (µM) | IC50 VE-822 (µM) | IC50 AZD7762 (µM) | IC50 AZD0156 (µM) | IC50 KU60019 (µM) |
|---|---|---|---|---|---|---|
| **Rh4** | 0.926 | 0.035 | 0.061 | 0.181 | 2.095 | 3.039 |
| **Rh30** | 0.480 | 0.035 | 0.082 | 0.217 | 1.533 | 7.667 |
| **Rh41** | 1.030 | 0.069 | 0.057 | 0.047 | 2.307 | 2.816 |
| **Kym1** | 0.811 | 0.069 | 0.208 | 0.230 | 3.599 | 7.189 |
| **RD** | 3.066 | 0.143 | 0.170 | 0.473 | 2.110 | 2.345 |
| **Rh18** | 13.030 | 0.303 | 0.818 | 1.994 | 6.225 | 11.260 |
| **T174** | 1.811 | 0.062 | 0.074 | 0.355 | 2.726 | 0.420 |
| **TE381.T** | 7.003 | 0.325 | 0.274 | 0.360 | 3.496 | 5.497 |
| **TE441.T** | 0.592 | 0.016 | 0.094 | 0.231 | 1.634 | 5.383 |

**Extended Data Fig. 2. PAX3-FOXO1-expressing alveolar rhabdomyosarcoma cells are hypersensitive to pharmacological ATR pathway inhibition**. **(a)** Dose-response curves rhabdomyosarcoma cell lines treated with ATR inhibitor VE-822 (*n*=3). (**b)** AUC of the previous curves (*P*=0.178 for ARMS vs ERMS). **(c)** Dose-response curves rhabdomyosarcoma cell lines treated with CHK1 inhibitor AZD7762 (*n*=3). **(d)** AUC of the previous curves (*P*=0.279 for ARMS vs ERMS). **(e)** Dose-response curves rhabdomyosarcoma cell lines treated with ATM inhibitor KU60019 (*n*=3). (**f)** AUC of the previous curves (*P*=0.519 for ARMS vs ERMS, *P*= 0.117 for ARMS vs Ctrl). (**g)** Dose-response cell viability curves for ARMS cell lines (red) and ERMS cell lines (blue) treated with the ATM inhibitor AZD0156 (*n*=3). **(h)** AUC of the previous curves (*P*=0.311 for ARMS vs ERMS). **(i)** Summary of calculated IC50 from the dose-response curves (two-sided student's t-test; error bars represent standard error of the mean).

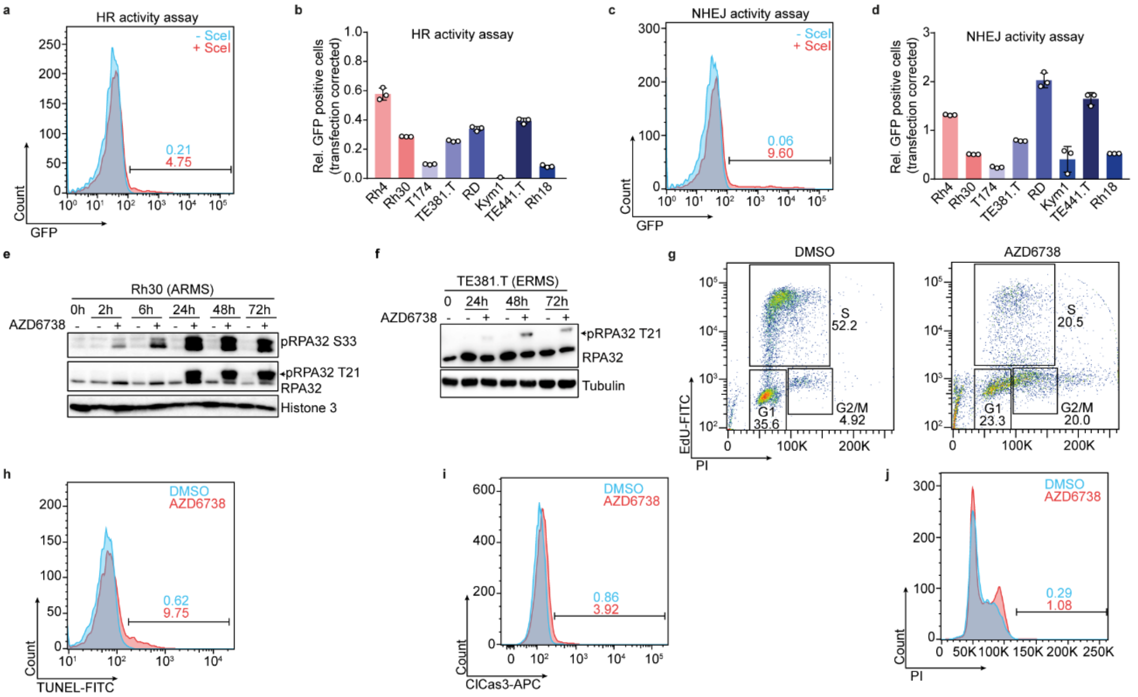


**Extended Data Fig. 3. Pharmacological ATR inhibition exacerbates replication stress and leads to genomic instability, apoptosis and cell cycle disruption. (a)** Representative histograms showing relative GFP signal in a rhabdomyosarcoma cell line co-transfected with a HR-GFP plasmid and a plasmid expressing SceI (red) or an empty backbone (blue) as measured using FACS. **(b)** Homologous recombination activity of several rhabdomyosarcoma cell lines normalized to a non-cutting control and corrected for transfection efficacy (*n*=3). **(c)** Representative histograms showing relative measure GFP signal in a rhabdomyosarcoma cell line co-transfected with a NHEJ-GFP plasmid and a plasmid expressing SceI (red) or an empty backbone (blue) as measured using FACS. **(d)** Non-homologous end joining activity of several rhabdomyosarcoma cell lines normalized to a non-cutting control and corrected for transfection efficacy (*n*=3; error bars represent standard error of the mean). **(e-f)** Western immunoblot of RPA32 phosphorylation in cells treated with ATR inhibitor AZD6738 (750 nM) over time for Rh30 (e) and TE381.T (f). **(g)** Representative histograms showing relative EdU and PI co-staining in the absence (left) and presence (right) of AZD6738 (750 nM) as measured using FACS. (**h)** Representative histograms showing relative TUNEL staining as measured using FACS. **(i)** Representative histograms showing relative cleaved Caspase3 staining as measured using FACS. **(j)** Representative histograms showing relative DNA content labeling (with propidium iodine, PI) as measured using FACS.

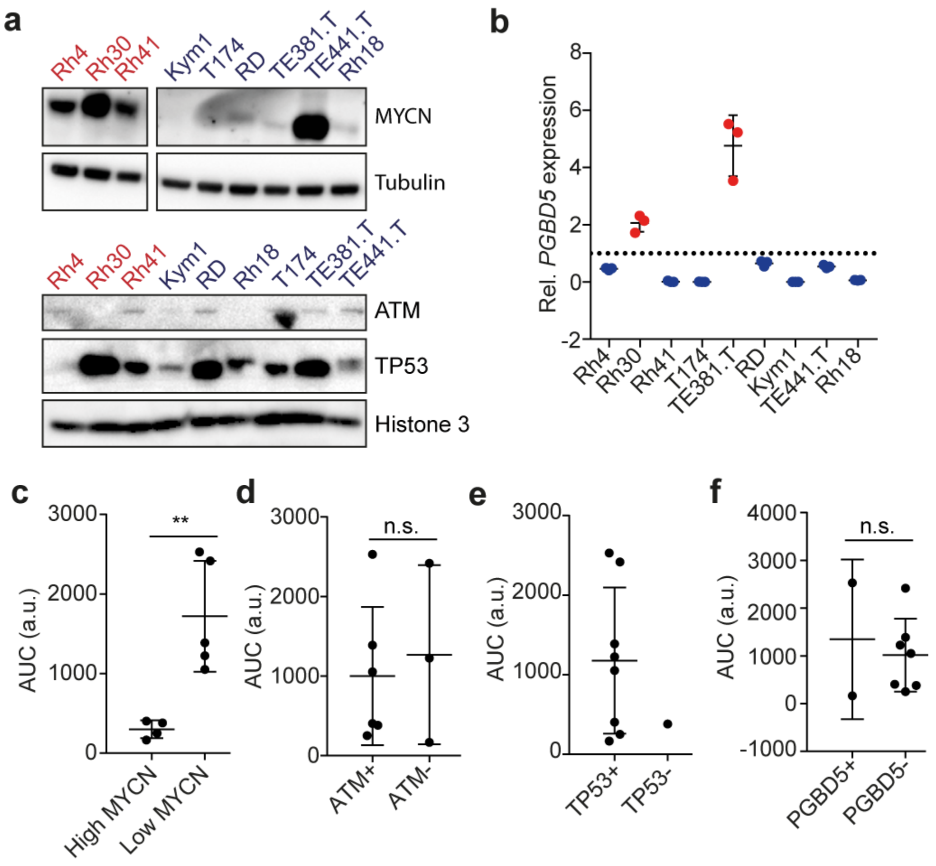


**Extended Data Fig. 4. Molecular factors associated with ATR inhibitor sensitivity in rhabdomyosarcoma cells**. (**a)** Western Immunoblot of MYCN, ATM and TP53 in rhabdomyosarcoma cell lines. **(b)** mRNA expression levels of *PGBD5* in rhabdomyosarcoma cell lines (*n*=3). **(c-f)** Comparison of AUC from dose response curves shown in Fig. 1b for MYCN (*P*=0.005; c), ATM (*P*=0.702; d), TP53 (e) and PGBD5 (*P*=0.677; f).

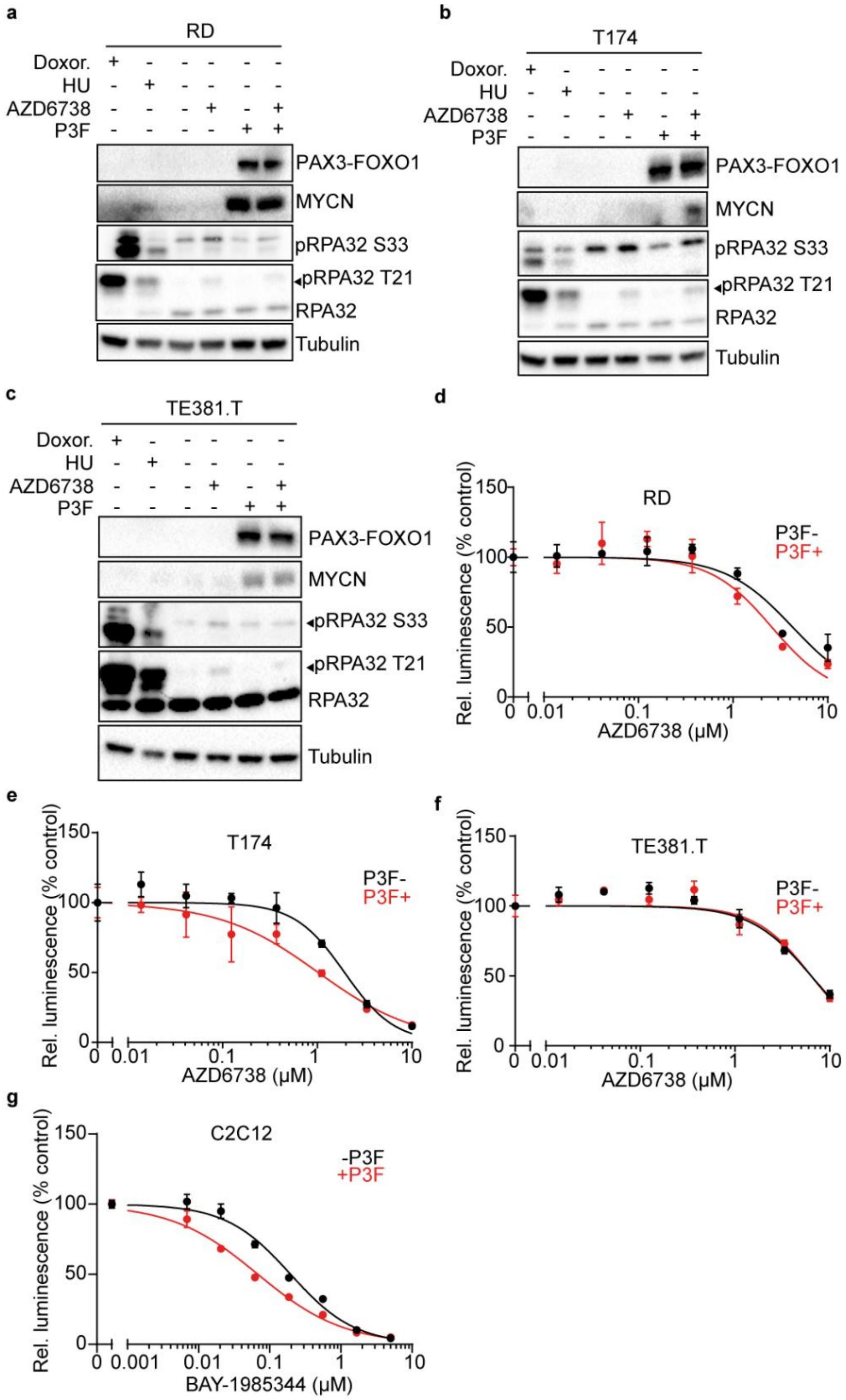


**Extended Data Fig. 5. Ectopic expression of PAX3-FOXO1 increases sensitivity to ATR inhibition in rhabdomyosarcoma cells. (a-c)** RD (a), T174 (b) and TE381.T (c) cells after induction of PAX3-FOXO1 expression with doxycycline (1000 ng/ml for 48 hours; 750 nM AZD6738 for 48 hours; doxorubicin and hydroxyurea treated cells, positive controls; Tubulin serves as a loading control). **(d-f)** Dose-response curves in RD (d), T174 (e) and TE381.T (f) cells treated with ATR inhibitor AZD6738 after doxycycline-induced ectopic expression of PAX3-FOXO1 (*n*=3). **(g)** Dose-response curves in C2C12 cells treated with ATR inhibitor BAY-1895344 after doxycycline-induced ectopic expression of PAX3-FOXO1 (*n*=3; error bars represent standard error of the mean).

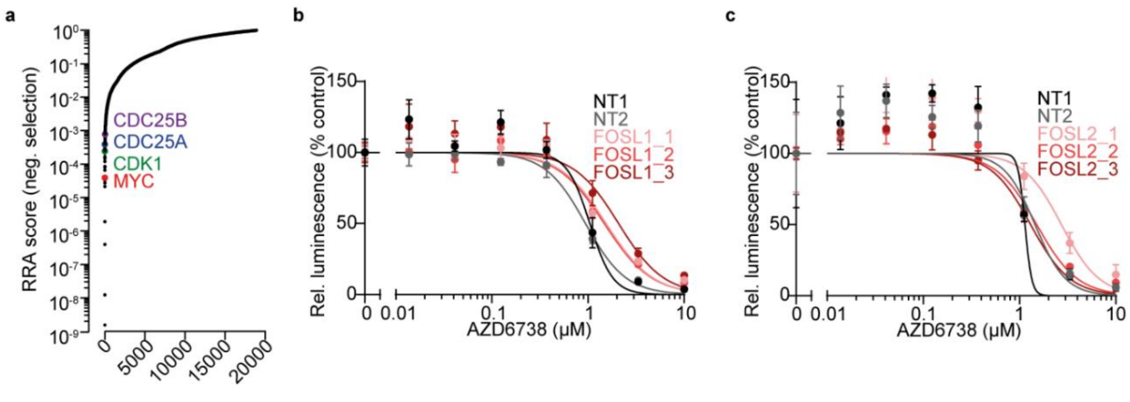


**Extended Data Fig. 6. *FOSB*, *FOSL1* and *FOSL2* expression induces resistance to ATR inhibition in rhabdomyosarcoma cells. (a)** Waterfall plot showing the negative robust rank aggregation (RRA) score of sgRNAs in Rh4 cells incubated in the presence of AZD6738 for 9 days compared to DMSO treated cells as analyzed using MaGECK. **(b-c)** Relative cell viability of cells stably expressing dCas9, lentiMPH and sgRNAs targeting *FOSL1* (b) and *FOSL2* (c) in the presence and absence of AZD6738 ($n$=3; two-sided student's t-test; error bars represent standard error of the mean).

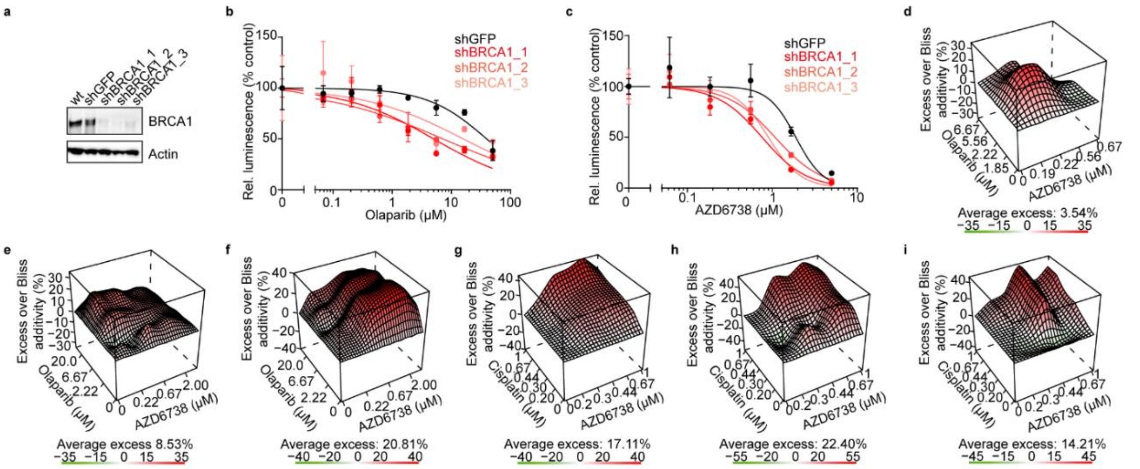


**Extended Data Fig. 7. ATR inhibition synergizes with cisplatin and olaparib in rhabdomyosarcoma cell lines. (a)** Western immunoblot of BRCA1 in Rh4 cells of stably expressing shRNAs targeting BRCA1 (shRNA targeting GFP was used as a control). **(b)** Dose-response curves for Rh4 stably expressing different shRNAs targeting BRCA1 and treated with PARP inhibitor olaparib (shRNA targeting GFP was used as a control) (*n*=3). **(c)** Dose-response curves for Rh4 transduced with different shRNA targeting BRCA1 treated with ATR inhibitor AZD6738 (shRNA targeting GFP was used as a control) (*n*=3; error bars represent standard error of the mean). (**d-f)** Excess over Bliss analysis of combined treatment with olaparib and AZD6738 in Rh30 (d), T174 (e) and TE381.T (f) cells (*n*=3). (**g-i)** Excess over Bliss analysis of combined treatment with cisplatin and AZD6738 in Rh30 (g), T174 (h) and TE381.T (i) cells (*n*=3).

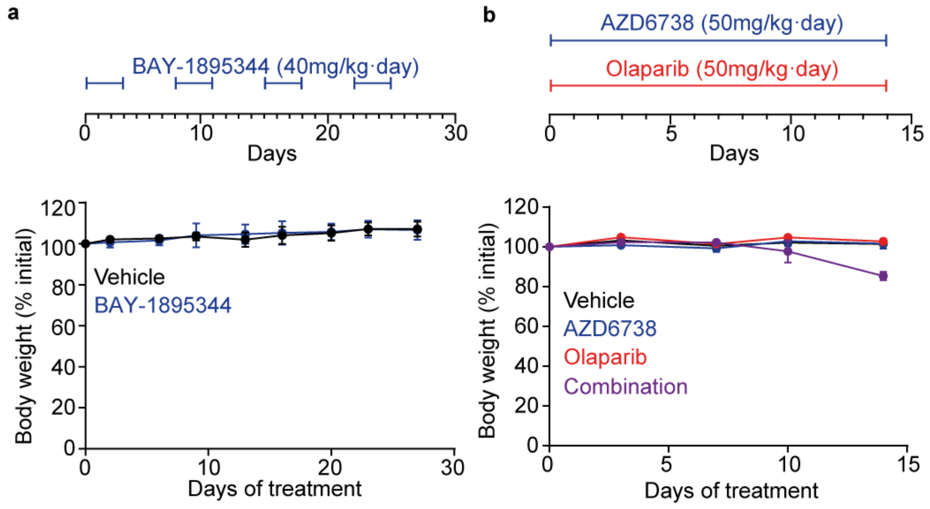


**Extended Data Fig. 8. Combined treatment of mice with AZD6738 and olaparib has no remarkable toxicity in mice harboring alveolar rhabdomyosarcoma PDX models.** **(a)** Timeline of the drug schedule. BAY-1895344 was given twice daily by oral gavage at 40mg/kg body weight per day on a 3 days on/4 days off schedule. Body weight over time of mice harboring the rhabdomyosarcoma PDX model subcutaneously xenografted and treated with BAY1895344 or a vehicle control. (n=7). **(b)** Timeline of the drug schedule. AZD6738 and olaparib are given by oral gavage at 50 mg/kg body weight per day. Body weight over time of mice harboring the rhabdomyosarcoma PDX model subcutaneously xenografted and treated with AZD6738, olaparib, both or a vehicle control (n=4; error bars represent standard deviations of 4 individual xenografted mice per group).

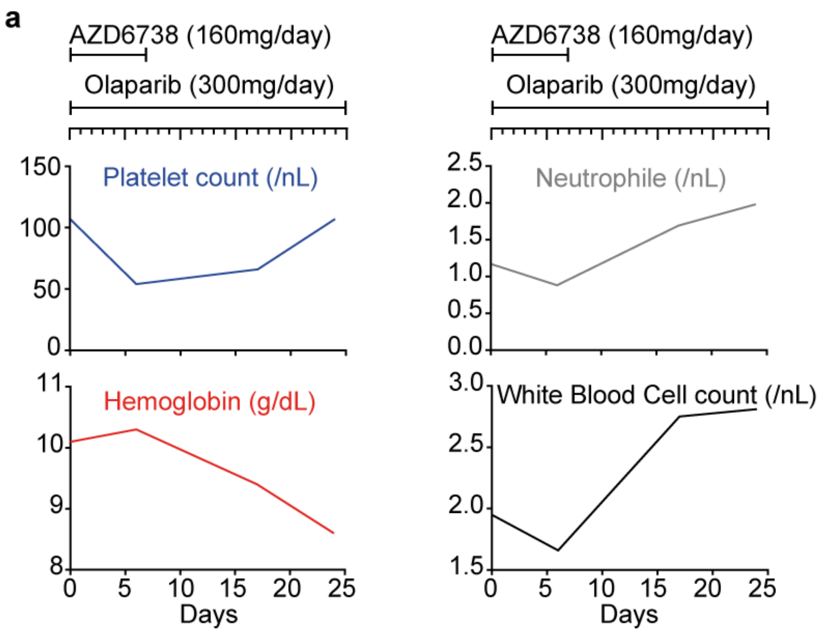


**Extended Data Fig. 9. Compassionate use of AZD6738 and olaparib in a patient suffering from relapsed metastasized PAX3-FOXO1-expressing alveolar rhabdomyosarcomas. (a)** Patient blood counts over the course of treatment with AZD6738 and olaparib.

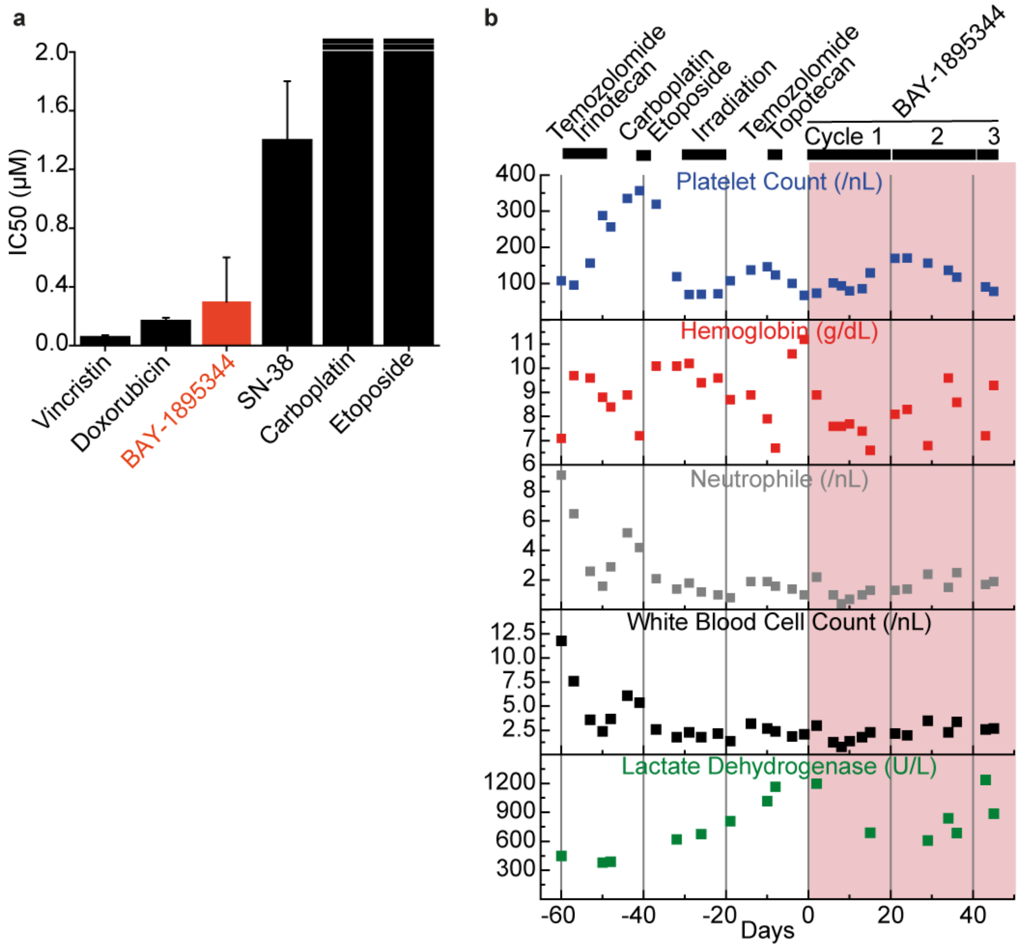


**Extended Data Fig. 10. Compassionate use of BAY-1895344 in a patient suffering from relapsed metastasized PAX3-FOXO1-expressing alveolar rhabdomyosarcomas. (a)** Half maximal inhibitory concentration ($IC_{50}$) for the anti-tumor effects of drugs against patient-derived rhabdomyosarcoma cells ($n$=3). Error bars represent standard error of the mean**. (b)** Key laboratory values in the second case report patient treated with BAY 1895344.

**Supplementary Table 1. Oligonucleotide primers**.

| Primer | Sequence (5’-3’) |
|---|---|
| **FOSB_Fwd** | GTG AGA GAT TTG CCG GGC TC |
| **FOSB_Rv** | AGA GAG AAG CCG TCA GGT TG |
| **FOSL1_Fwd** | GCC CAC TGT TTC TCT TGA GC |
| **FOSL1_Rv** | GAT GGA GAG TGT GGC AGT GA |
| **FOSL2_Fwd** | GCC CAG TGT GCA AGA TTA GC |
| **FOSL2_Rv** | GGG CTC CTG TTT CAC CAC TA |
| **HPRT1_Fwd** | TGA CAC TGG CAA AAC AAT GCA |
| **HPRT1_Rv** | GGT CCT TTT CAC CAG CAA GCT |
| **PGBD5_Fwd** | CAG CCT CTG GGT CAG ACA AT |
| **PGBD5_Rv** | GCT TAT TCT TCA GCG CAT CC |

**Supplementary Table 2. Antibodies.**

| Antibody | Company | Catalog number |
|---|---|---|
| **FOXO1 (And PAX3-FOXO1)** | Santa Cruz Biotechnology | sc-374427 |
| **Tubulin** | Cell Signaling Technology | 3873 |
| **Actin** | Cell Signaling Technology | 3700 |
| **Histone 3** | Cell Signaling Technology | 4499 |
| **RPA32 T21** | Abcam | ab61065 |
| **RPA32 S33** | Bethyl Laboratories | A300-246A |
| **MYCN** | Santa Cruz Biotechnology | sc-53993 |
| **ATR** | Cell Signaling Technology | 13934 |
| **ATM** | Cell Signaling Technology | 92356 |
| **CHK1** | Cell Signaling Technology | 2360 |
| **CHK2** | Cell Signaling Technology | 2662 |
| **TP53** | Santa Cruz Biotechnology | sc-98 |
| **BRCA1** | Merck Millipore | OP92-100UG |
| **BRCA1 S1524** | Bethyl Laboratories | A300-001A |
